# Evolutionary Emergence of First Animal Organisms Triggered by Environmental Mechano-Biochemical Marine Stimulation

**DOI:** 10.1101/2020.12.03.407668

**Authors:** Ngoc Minh Nguyen, Tatiana Merle, Florence Broders, Anne-Christine Brunet, Florian Sarron, Aditya Jha, Jean-Luc Genisson, Eric Rottinger, Emmanuel Farge

## Abstract

The evolutionary emergence of the first animals is thought to have been intimately associated to the formation of a primitive endomesodermal gut (*i.e* gastrulation) from ancestral multi-cellular spheres, blastulae, more than 700 million years ago. However, the biochemical cues having been at the origin of endomesoderm formation remain a mystery.

Here we find that hydrodynamic mechanical strains developed by sea wavelets on pre-bilaterian *Nematostella vectensis* and pre-metazoan *Choanoeca flexa* representatives, which common ancestor dates back to more than 700 million years ago, can trigger gastrulation in a Myo-II dependent mechanotransductive process. Gastrulation in turn induces endomesoderm first biochemical specification through the mechanical activation of the βcat pathway in pre-bilaterian *Nematostella vectensis*, like in Drosophila and zebrafish embryos, which common ancestor dates back to 600-700 million years ago.

These observations converge to animal emergence that has been mechanotransductively triggered by wavelet mechanical strains on the sea-shore in multicellular choanoflagellates through Myo-II more than 700 million years ago, a process achieved in first metazoan through mechanosensitive Y654-containing βcat evolutionary emergence found as conserved in all metazoan.

**One sentence summary:** Marine hydrodynamic strains have activated first gastric organ formation from ancestral pre-animal cell colonies.

---

The appearance of the first animal organisms is thought to have been triggered by the evolutionary emergence of the biomechanical morphogenetic invagination (gastrulation) and biochemical specification of the endomesoderm (EM) from blastulae primitive multi-cellular tissues, that led to the first primitive digestive organ (*1-4*). However, due to the nonconserved biochemical signals involved in today’s metazoan early embryos, in both the molecular Myosin-II (Myo-II) activation leading to biomechanical gastrulation (*5-8*) and the βcat signalling pathway activation leading to EM versus ectoderm biochemical first tissue specification (*9*), the evolutionary origins of first multicellular animal emergence remain puzzling.

Interestingly, the activation of Myo-II, and the activation of the β-catenin pathway via phosphorylation of Y654-βcatenin, have been found to be mechanotransductively triggered in leading to EM morphogenesis and specification in gastrulating species (*Drosophila* and Zebrafish) that diverged from the first bilaterian animals around 570 million years ago (*9-11*). Thus, mechanotransduction seems to be a conserved feature to trigger intracellular biochemical processes leading to EM gastrulation and specification. This further suggests that mechanotransduction might also be an ancestral property inherited from pre-bilaterian animals or even pre-metazoan “multi-cellular” spheres, preceding the emergence of metazoans.
To test this hypothesis within the mechanical environmental context of most ancient animals, and track the ancestrally and inheritance lineage of EM mechanical induction, we applied soft hydrodynamic mechanical strains developed by sea shore wavelets on multi-cellular blastulae of either the pre-bilaterian cnidarian *Nematostella vectensis (Nv)* and the pre-metazoan choanoflagellate *Choanoeca flexa*, from which bilaterians appear to have diverged 600-700 million and more than 700 million years ago, respectively (*12, 13*).

## Mimicking sea-shore wavelets triggers cnidarian *Nematostella* blastulae embryo gastrulation

In most metazoa, gastrulation consists in the first morphogenetic movement of embryogenesis and is thought to be the fossil structure of the first animal ancestral digestive track morphology (*1, 2*). As shown in Fig. 1A, the *Nv* embryo, still isotropic at 18 hours post fertilization (h), presents after 21h an invagination (of depth *d*) together with an apex constriction known to drive the invagination (orange arrow), as seen on actin labelled gastrulating embryos. In today’s Drosophila bilaterian early embryos, internal and external mechanical strains were found to mechanotransductively trigger active movements of gastrulation, thereby initiating embryonic biomechanical morphogenesis (*11, 14-16*). We wonder here, in an evolutionary perspective, whether such mechanism could have been inherited from earliest animals found in the sea close to the shore, with gastrulation stimulated by the marine hydrodynamic environment of the seashore.

**Figure 1:**
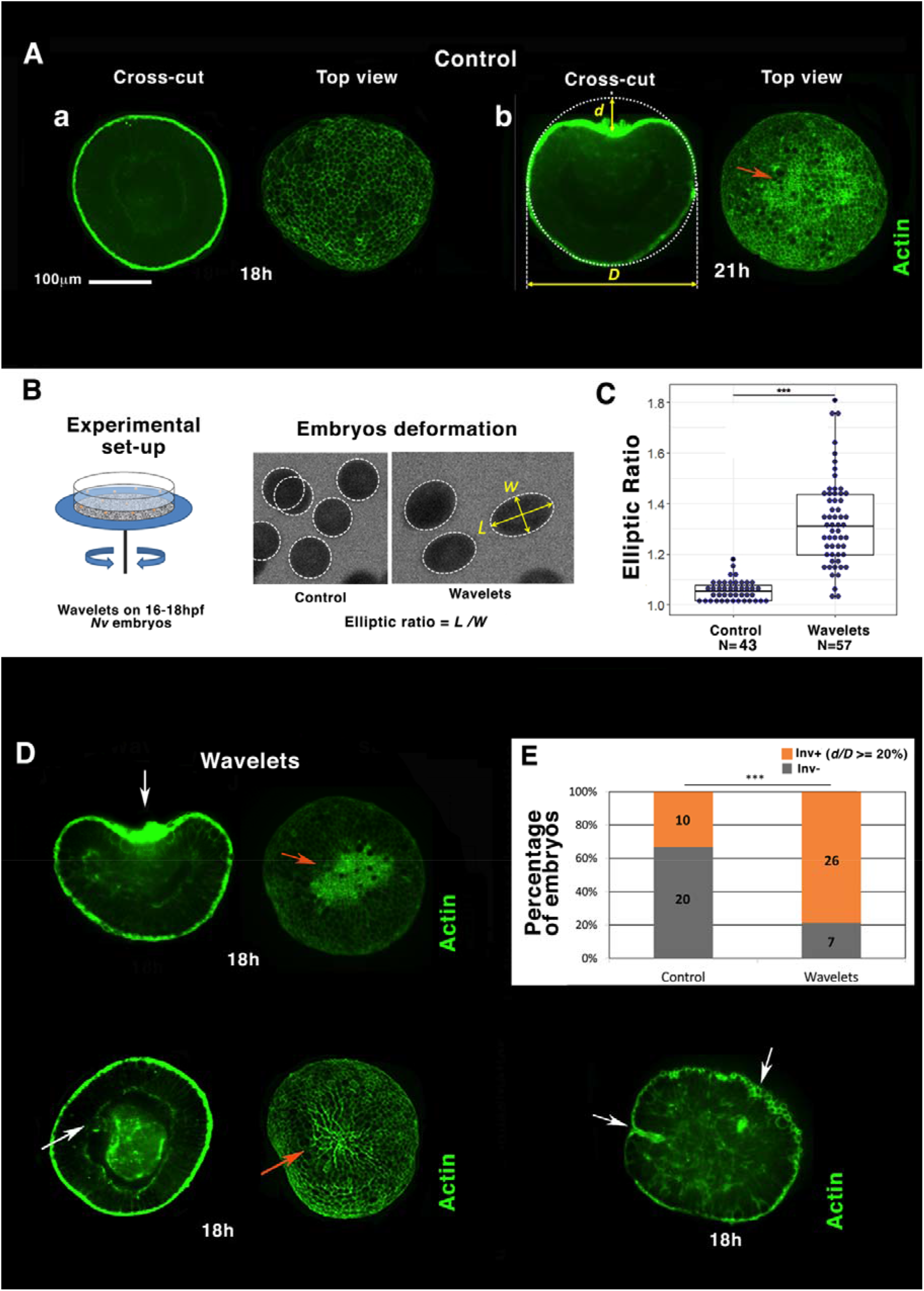
Hydrodynamic stimulation of *Nematostella* embryos gastrulation by marine wavelets. **A** Most representative gastrulation state population of embryos at **a** 18hpf and **b** 21hpf. **B** Setup: mimicking sea ocean wavelets on the shore by 2.5s period shaking of a petri dish with sand on its ground. Hydrodynamic deformation of jelled embryos with sand, induced by wavelets. **C** Elliptic deformation of embryos quantification. Mann-Whitney statistical test. **D** Embryos representative of the majority of individuals submitted to wavelets. **E** Quantitative characterisation of the percentage of gastrulating embryos (in orange; embryos number inside columns), submitted to wavelets compared to control, based on significant normalized invagination depth *d/D* measurement higher or equal to 20% (see text, Fisher statistical test). N=2 for each experiment.

To that end, we began by subjecting cnidarian *Nv* blastulae early embryos to the characteristic soft wavelets of a seashore, a way to initiate the testing of the hypothesis on the 600-700 millions years ago common ancestor of bilaterian and cnidarian embryos. As sketched in Fig. 1B, a petri dish containing the embryos was shaken with a rotation amplitude of 2cm radius, a speed of 85 rounds-per-minute and an rotation orientation inverted any 2.5 s – corresponding to a flow with a velocity of ~10cm/s and a periodicity of ~2s typical of a seashore environment. Embryos were stimulated within physiological conditions, *i.e* inside their deformable protective jelly (*17*). Sand (~ 1mm size, Fig. S1A) was first glued on the petri dish to approach physiological conditions that enhance the friction at the jelly surface on the ground. Taking advantage of the transparency of the jelly, embryos were observed by fast imaging and found to be deformed by the substrate rotation, so as to adopt an ellipsoidal shape characterized by its elliptic ratio (Fig. 1B, see Materials and methods). Mean elliptic deformation was found to increase from the nearly 5% under static conditions to 30% under hydrodynamic flow (Fig. 1C, Fig. S1B).

Jellied blastulae were stimulated with wavelets for 2 hours between 16h and 18h, that is, in a stage known normally to be non-gastrulating. Embryos were then dejellied and fixed and actin was labelled with phalloïdin at 18h stage (3h before the time of gastrulation of un-perturbed embryos, Fig. 1A). As seen in Fig. 1D, E-wavelets, and in contrast to Fig. 1Aa,E-control, we observed that embryos gastrulation was initiated at 18h in the majority of hydrodynamically stimulated embryos, with no obvious link between the symmetry of the stimulating deformation (seen in Fig. 1B) and the symmetry of the responding gastrulation. The gastrulating embryos (with diameter *D*) subject to wavelets show an invagination (with depth *d*, white arrow) with a normalized depth *d/D* equal or larger than 20% (with a precision measurement of 5%) and accompanied by an apex constriction in the invaginating tissue (Fig. 1D, orange arrow). This corresponds to a change of curvature from +10mm^-1^ to −8 mm^-1^, where the change of sign evidences the emergence of the invagination, and the absolute value its development.

In ~ 60% of cases, gastrulating embryos showed anomalous shapes, with large invaginations enlarging the lateral size of the embryo (first row of Fig.1D, ~ 30% of cases), and tubular-like and possibly multiple invaginations (last photo, second row of Fig.1D, ~ 30% of cases), also satisfying the criterion *d/D* > 20%. These cases are thus added to the global statistics presented in Fig. 1E where almost 80% of the embryos exhibit an invagination after 18h if subjected to the wavelets, instead of 33% in the control situation. Despite the fact that the majority of stimulated invaginations look anomalous (wider, tube-like, multi invagination sites, etc...), all embryos for which we let enough time to reach maturity were found to be as viable as control embryos. Interestingly, dejellied embryos also showed a significant gastrulating response in wavelets, even though less sensitive. Hence hydrodynamic strain stimulation directly applied on embryos initiates gastrulation as well (Fig. S1C and Sup info1). This is an interesting observation in the context of evolution where embryos of the last common ancestor of bilaterian and cnidarian may have been simply drifted in water.

In addition to the fact that most of stimulated invaginations are anomalous in shape and number (Fig. 1D), the expression level of the developmentally expressed gene *bmp1-like*, known to be strongly up-regulate between 18h and 20h of development (NvERTx.4.130707 / NvERTx data base (18)), was not up-regulated in stimulated compared to control embryos (Fig. S2A,B). This ensured that hydrodynamically induced gastrulation was not a consequence of a global acceleration of development, for instance due to a putative increased oxygenation under shaking conditions. A quantified flow-meter hydrodynamic oscillating laminar flow of ~10cm/s within the same but standing still petri dish (Fig. S3A) confirmed the hydrodynamic stimulation of invaginations (Fig. S3B,C).

This shows that hydrodynamic mechanical strains characteristic of the sea-shore soft waves are inductive of embryonic gastrulation in the *Nv* cnidarian representative.

## Hydrodynamical stimulation of *Nematostella* gastrulation is Myo-II and oral-aboral polarity dependent

Apical constriction was observed in the invaginating tissue of gastrulating embryos with an inward curvature of at least 20% invagination depth (Fig. 1A-E). Since apical constriction leading to gastrulation is Myo-II dependent in many species (*19-21*), we tested in physiological jellied embryos conditions whether mechanical hydrodynamic stimulation of gastrulation requires Myo-II. Using the ML7 inhibitor of Myo-II (*22*), we observed the inhibition by ML7 of hydrodynamically induced apical constriction and gastrulation. At 18h, the percentage of invaginating embryos subjected to wavelets then reduces by a factor of ~3 and recovers the non-stimulated control percentage (Fig.2A,B, Sup info *2*). This result shows that Myo-II motor activity is required for the underlying mechanotransductive process of hydrodynamically induced gastrulation by wavelets.

**Figure 2:**
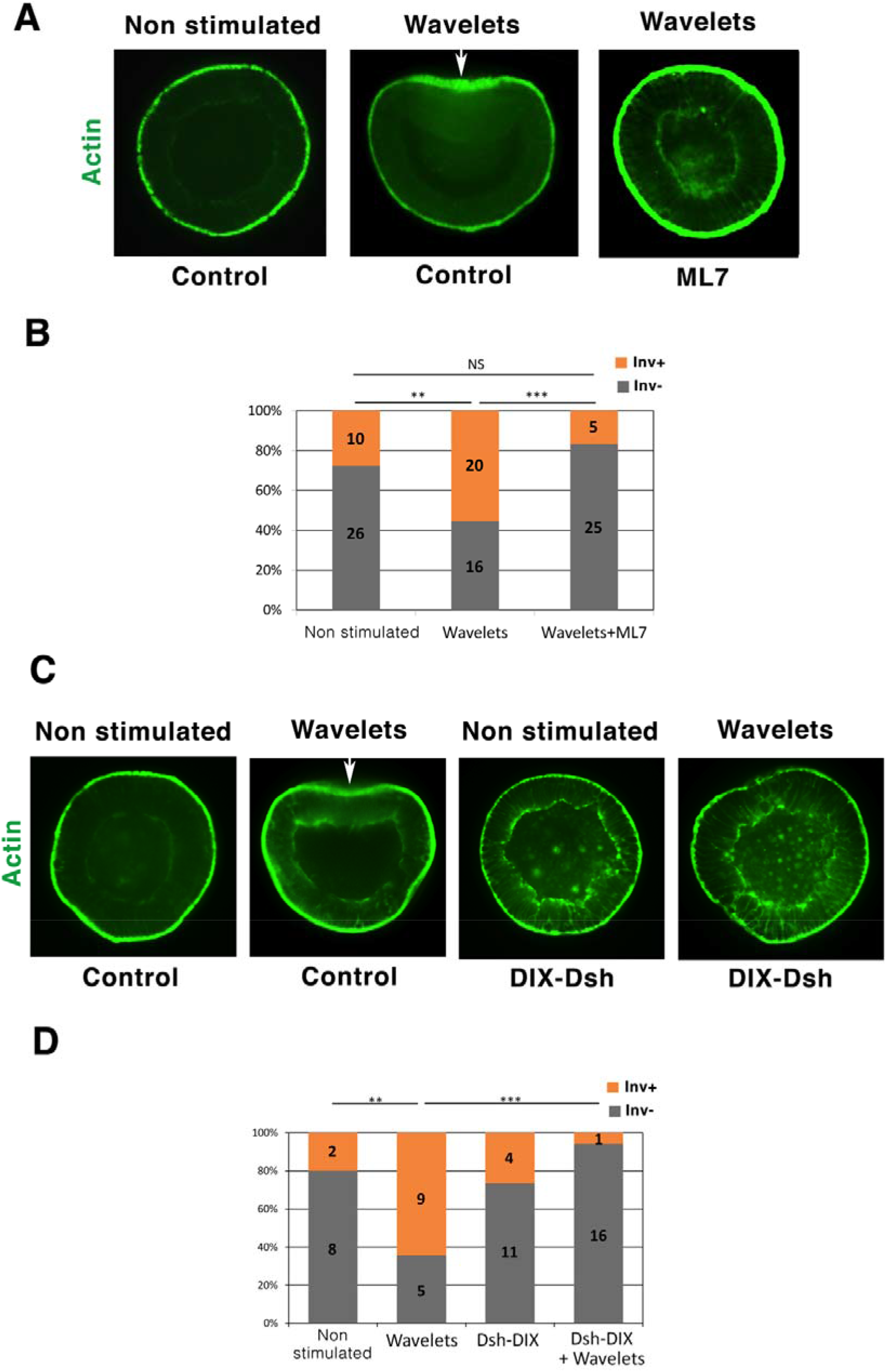
Hydrodynamic stimulation of *Nematostella* embryos invaginations is Myo-II and Dsh dependent. **A** 18hpf hydrodynamically stimulated *Nematostella* embryos in the presence of the ML7 Myo-II inhibitor. Control: ethanol, vehicle of ML7 (see methods). ML7 treated embryos show a slightly higher thickness than controls. **B** Quantitative analysis. Statistical test: Fisher. **C** 18hpf hydrodynamically stimulated *Nematostella* embryos injected with the Dsh-DIX-ΔN dominant negative. Control: water injected. D Quantitative analysis. Control embryos with wavelets shows 60% invagination in contrast of the 2 other groups in which this percentage is divided by a factor of at least 3. Some residues are observed inside the embryo, inherent to Dsh-DIX injection *per se*. Statistical test: Fisher. N=2 for each experiment.

In addition, if hydrodynamically induced invaginations could interestingly be found in multiple sites in a few cases (~25%, Fig. 1F-right), these invaginations remain grouped and polarized in one hemisphere of the embryo. This observation suggests the existence of a Myo-II response that is dependent on embryonic oral-aboral pre-patterning, known to be initiated by the oral expression of Dishevelled (Dsh) (23). To confirm this fact, we compared embryos defective in Dsh due to DIX-Dsh injection (23) with control embryos, with and without wavelets. The statistics again shows 60% of wavelets stimulated gastrulating embryos (instead of 20% for control embryos), most entirely inhibited in Dsh defective embryos injected with DIX-Dsh (Fig. 2C,D and Sup info 3), also known to be gastrulation defective at 21h in the absence of stimulation (23).

This indicates the existence of a Dsh-dependent pre-patterned mechanosensitive pathway leading to Myo-II-dependent hydrodynamically induced activation of gastrulation in *Nv*.

## Hydrodynamically induced gastrulation triggers *fz10* EM specification gene in *Nematostella* embryos

In bilaterian early embryos, gastrulation (invagination in Drosophila) and first morphogenetic movements of embryogenesis (epiboly in zebrafish) lead to the mechanical activation of endomesodermal gene expression via the activation of the β-catenin pathway (*9, 24*). Latter is mediated by the mechanical stimulation-induced phosphorylation of the β-catenin residue Y654 (pY654-βcat). We thus tested in *Nv* whether such property was already present in the common ancestor of bilateria and cnidaria.

In line with that idea, we first observed by *in situ* hybridation labelling, that expression of the endomesodermal β-catenin target *gene fz10* (*17, 25*) correlates with EM invagination in *Nv* at gastrulation stage (Fig. 3A,B). With embryo top views showing purple *fz10* labelling inside embryos into gastrulating tissues. Similarly, we found that hydrodynamically stimulating gastrulation in 18hpf *Nv* embryos (Fig. 1F) triggered *fz10* expression at the gastrulation site (Fig.3C,E). Interestingly, the percentage of *fz10* positive individuals of both the control and wavelets stimulated embryos correlates strongly with the percentage of gastrulating positive individuals (Fig. 1E). In contrast, hydrodynamically stimulated embryos in which gastrulation was blocked by the Myo-II inhibitor ML7, did not express *fz10* (Fig.3D,F). These results indicate that hydrodynamic strains *per se* could not stimulate *fz10* expression. Rather, the mechanical strains generated by gastrulation are the cause of the mechanotransductive and Myo-II dependent expression of the endomesodermal *gene fz10* in the invaginating *Nv* tissue.

**Figure 3:**
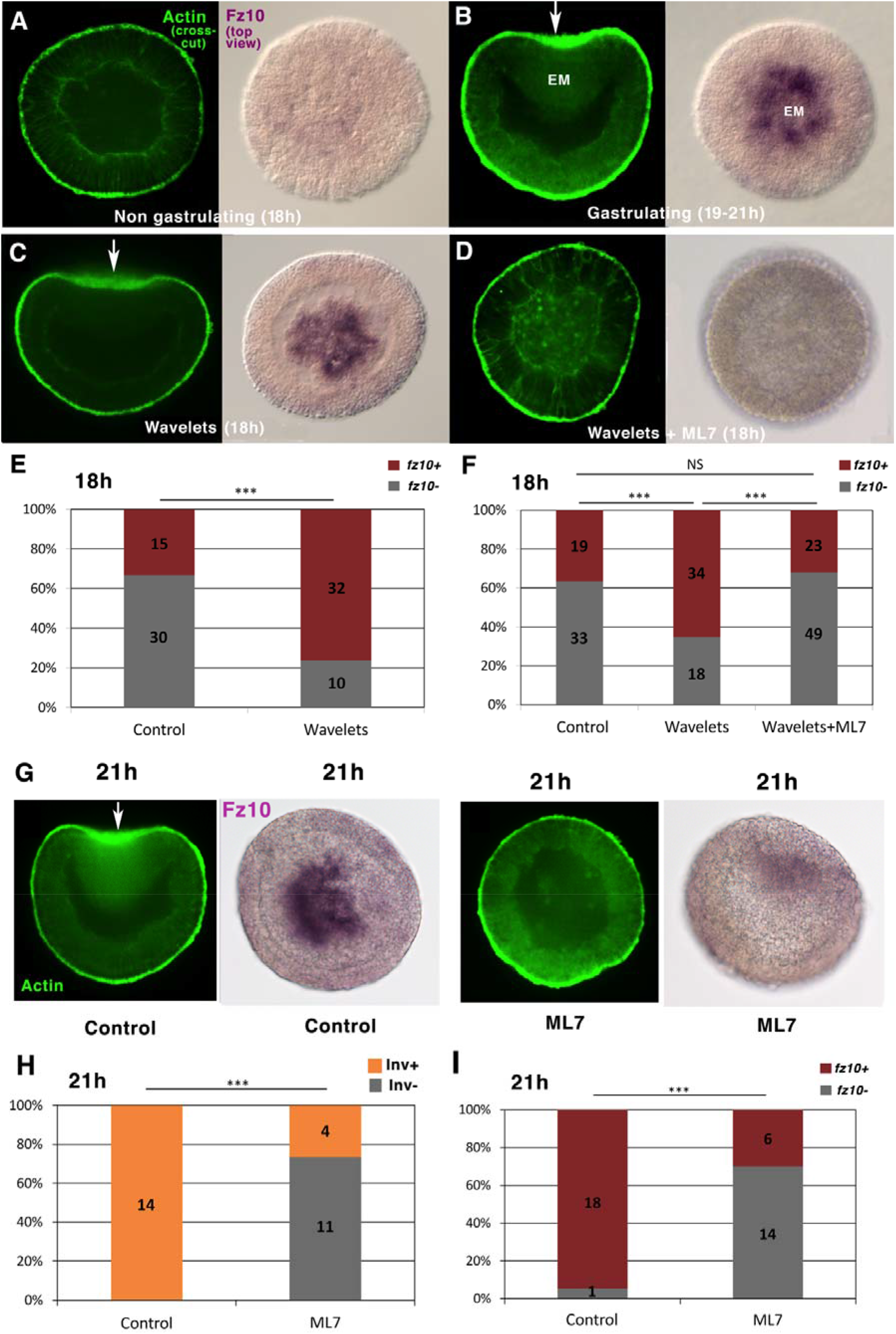
Hydrodynamically induced and endogenous gastrulation of *Nematostella* embryos mechanotransductively stimulates the endomesodermal *fz10* gene expression in a Myo-II dependent process. **A** *fz10* expression in 18hpf non-invaginating embryos, **B** and in the invaginating endomesoderm of 21hpf embryos. **C** *fz10* expression in the invaginating tissue of 18hpf hydrodynamically stimulated embryos, D and in hydrodynamically stimulated non-gastrulating embryos treated with ML7. **E,F** Quantitative analysis, with an increase of a factor of 3 of *fz10* individuals in the presence of wavelets only in E. Control of F: ethanol, vehicle of ML7. **G** *fz10* expression in non-gastrulating embryos ML7-treated 21hpf embryos, **H,I** and quantitative analysis. Control: ethanol, vehicle of ML7. Statistical test: Fisher. N=2 for each experiment.

Consistently, in non-stimulated embryos, *fz10* expression was impaired in ML7-treated non-gastrulating embryos compared to 21hpf gastrulating embryos (Fig.3G,H,I), additionally showing that endogenous (non-mechanically induced) *Nv* gastrulation mechanotransductively stimulates *fz10* EM gene expression, like in *Drosophila* and zebrafish embryos (*9, 24*).

This shows the *fz10* EM specification gene expression is mechanically induced in response to hydrodynamically stimulated or endogenous *Nv* gastrulation.

## Hydrodynamically induced gastrulation triggers *Nematostella* embryos EM specification in a Y654-β-cat phosphorylation dependent process

We then checked the behaviour of the mechanosensitive β-cat that targets *fz10*. We found that Y654-βcat phosphorylation, that leads to the release of β-cat from the junctions to the cytoplasm and nucleus in bilaterian embryos at gastrulation and epiboly (*9, 24*), is induced at Nv gastrulation, into the invaginating EM in 21h embryos (white arrows), and is inhibited in ML7 treated 21hpf embryos lacking gastrulation (Fig. 4A,B, Fig. S4 with Sup Info 4,5). pY654-βcat oral-aboral gradient was found to be on the order of 20%, which is in line with the EM-ectoderm gradient of Y654-βcat mechanically phosphorylated in the invaginating and stretched EM of gastrulating Drosophila and epibolying zebrafish embryos, respectively (9) (Fig. 4A,B).

**Figure 4:**
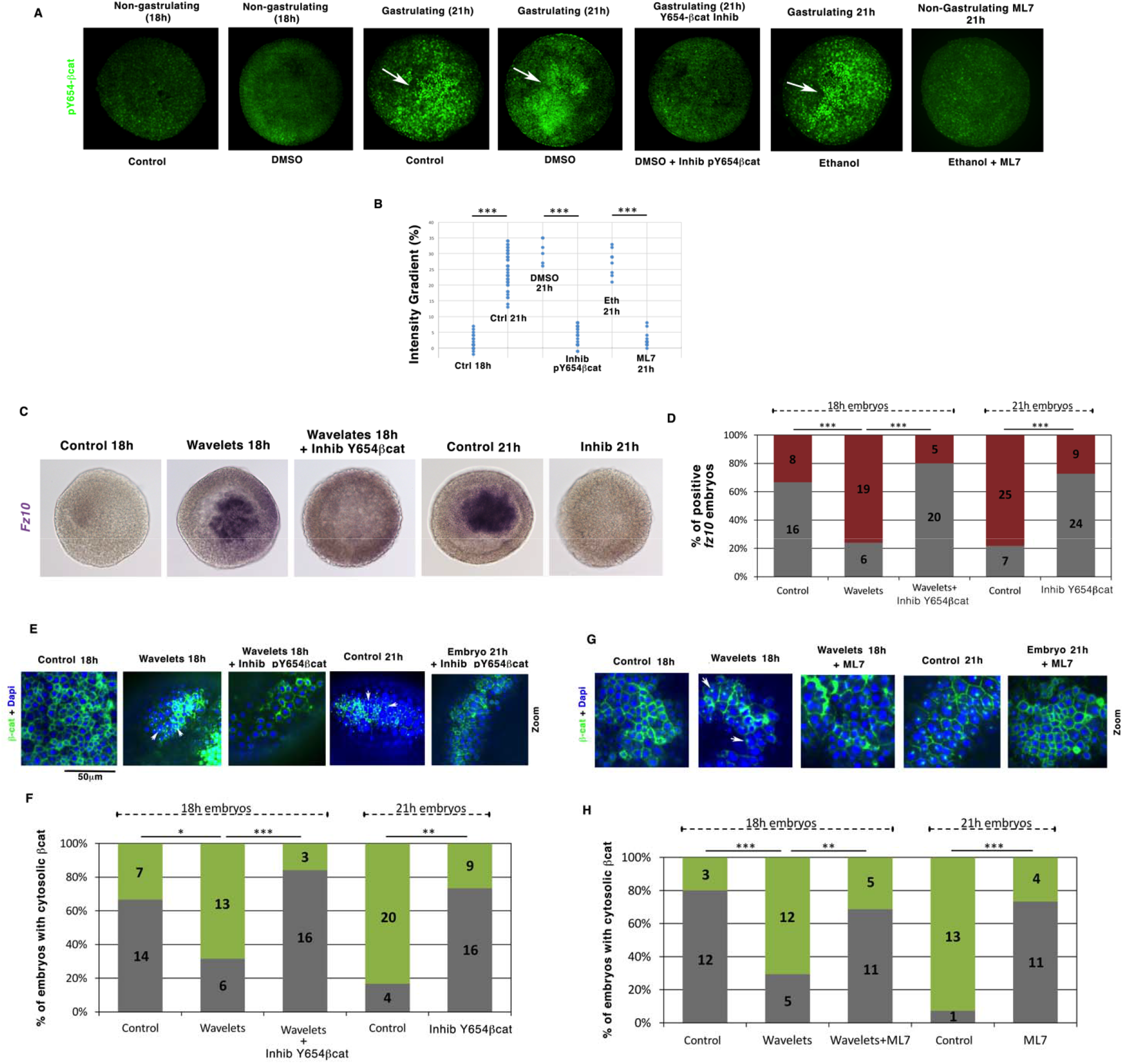
Mechanical induction of the endomesodermal *fz10* gene expression is initiated by the mechanical activation of Y654-β-cat phosphorylation. **A** Y654-β-cat phosphorylation in blastulae (18hpf), gastrulae (21hpf) and in 21hpf gastrulae embryos treated, with either InhibY654-βcat, the inhibitor of Y654 phosphorylation or ML7, the inhibitor of gastrulation. **B** Quantification of the maximal gradient of A. **C** Expression of *fz10* in 18h embryos stimulated with wavelets and 21h gastrulae embryos, treated with InhibY654-βcat. **D** Quantification of C. **E** Nuclear β-cat translocation state in the non-invaginating blastulae 18hpf tissue, in the 21hpf invaginating tissue of the gastrulating embryos, and in the hydrodynamically stimulated 18hpf and 21hpf invaginating tissues of gastrulating embryos treated with the Y654-β-cat phosphorylation inhibitor. Control and wavelets: DMSO, vehicle of Inhib-Y654-β-cat. **F** Quantification of E. **G** Nuclear βcat translocation states in both non-gastrulating ML7-treated 18h hydrodynamically stimulated and 21h embryos. Control and wavelets: Ethanol, vehicle of ML7. H Quantification of G. Statistical tests: Fisher for embryo number counting and Mann-Whitney for quantitative data by embryo. N=2 for each experiment.

This shows Y654-βcat phosphorylation as mechanically induced by the driving forces of gastrulation morphogenetic movements.

Blocking Y654-βcat phosphorylation with a specific inhibitor from 18h to 21h (Fig. 4A,B, Fig. S4 see Materials and Methods) prevented *fz10* expression in 18h embryos in which gastrulation was hydrodynamically stimulated during two hours, as well as in 21h gastrulating embryos (Fig. 4C,D and Sup info 6), This shows that the mechanical activation of the EM gene *fz10* by the hydrodynamically induced and endogenous morphogenetic movement of gastrulation requires the pY654-βcat mechanically induced by gastrulation movements.

Consistently, βcat cytosolic (cytoplasmic and nuclear) translocation, not observed in the non-gastrulating 18hpf embryos, was observed in 21h gastrulating embryos (Fig. 4E,F and Sup Info 7). Moreover, blocking Y654-βcat phosphorylation with the specific inhibitor from 18h to 21h (Fig. 4A,B) prevented βcat cytosolic (cytoplasmic and nuclear) translocation in 18h embryos in which gastrulation was hydrodynamically stimulated from 16hpf to 18hpf, as well as in 21h gastrulating embryos (Fig. 4E,F).

In addition, βcat cytosolic translocation was defective in 16hpf-18hpf hydrodynamically stimulated embryos gastrulation defective embryos treated with the ML7 inhibitor of Myo-II and in 21hpf ML7 treated non-gastrulating embryos (Fig. 4G,H).

This showed that mechanical induction of the cytoplasmic and nuclear translocation of βcat and of the expression of its *EM fz10* target gene by gastrulation morphogenetic movements are dependent of the mechanical stimulation of pY654 βcat by hydrodynamically induced, or endogenous gastrulation morphogenetic movements in the cnidarian *Nv*.

## Mimicking sea-shore wavelets triggers gastruiation-like inversion in the pre-metazoan multi-cellular choanoflagelatte Flexa

To test whether hydrodynamic stimulation of gastrulation dates back to organisms preceding the emergence of metazoa, we submitted the pre-metazoan multi-cellular choanoflagellate Flexa, to wavelets stimulation. Flexa has the ability to invert in response to light through an ancestral gastrulation-like process, as denoted by flagellate observed outward instead of inward, which consists in the primitive form of gastrulation in unclosed pre-metazoan spherical tissue (*26*). Here we inhibited the Flexa light sensor (see materials and methods). Strikingly, we found light-independent hydrodynamically induced stimulation of gastrulation-like inversion at 105 rpm, with flagellate outward after stimulation on Flexa (Fig. 5A,B,E).

**Figure 5:**
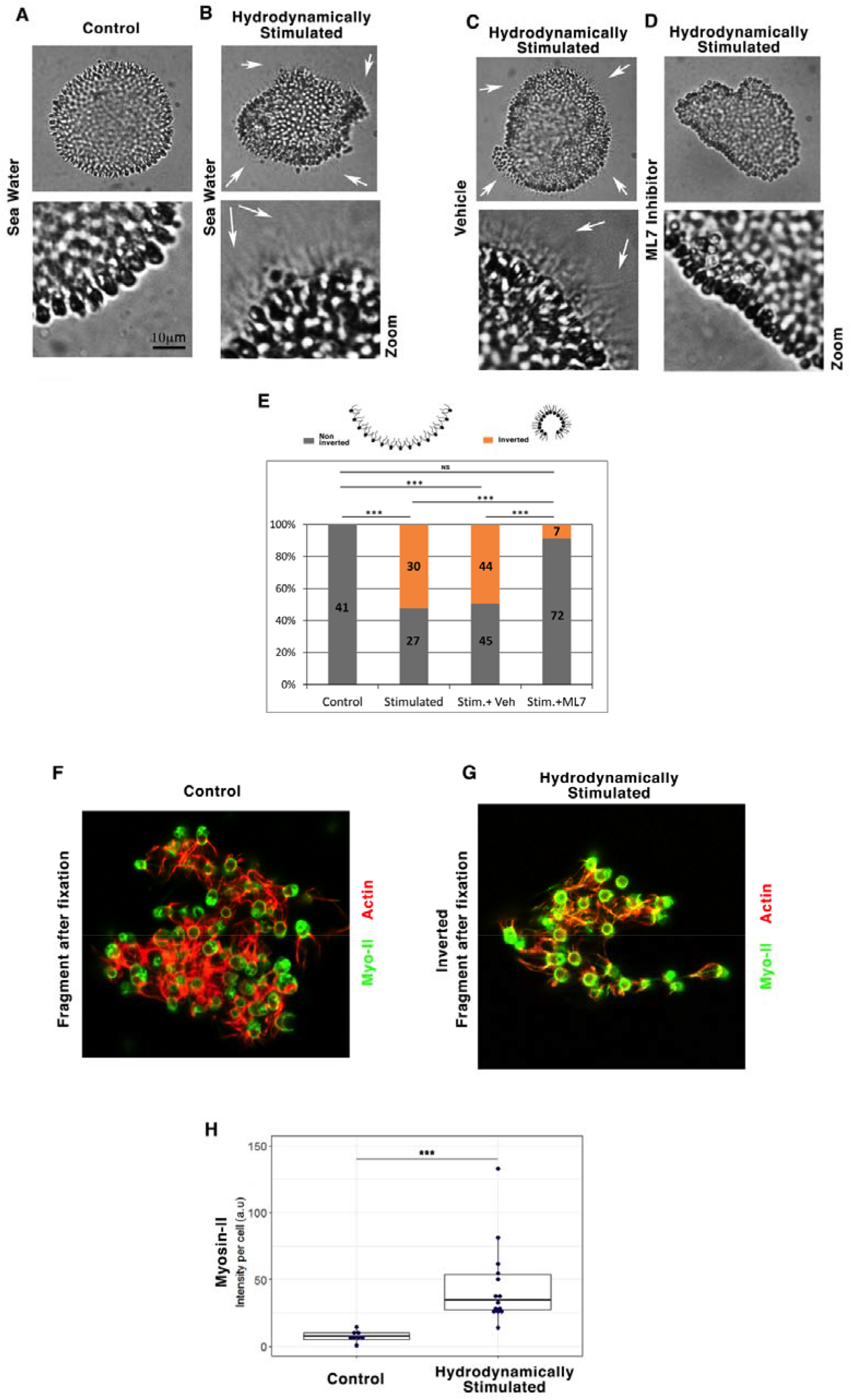
Myo-II dependent hydrodynamic stimulation of gastrulation-like inversion is conserved in Flexa choanoflagellates. **A** Multicellular Flexa choanoflagellate before hydrodynamic stimulation (no flagellate outside), **B** and after hydrodynamic stimulation (white arrows: flagellate outside). **C** Inversion status of hydrodynamically stimulated Flexa treated with the ethanol vehicle of ML7 alone (white arrows: flagellate outside), D and with the Myo-II inhibitor ML7 (no flagellate outside). Vehicle is Ethanol. **Zooms:** with enhanced contrast to check for flagellate presence. **E** Quantitative analysis. Scheme from (Brunet et al, 2019). **F** Myo-II and actin in Flexa before hydrodynamic stimulation, **G** and after hydrodynamic stimulation. **H** Quantitative analyses of Myo-II expression per cell 20 to 30 min after hydrodynamic stimulation initiation. Immunofluorescence representative of n=8 controls, and of n= 6 inverted structures of the hydrodynamically stimulated Flexa. N=2 experiments by case. Statistical tests are Fisher for histograms and Mann-Whitney for quantitative analysis.

Hydrodynamically induced inversion also triggered a transition to a swimming state possibly due to the freed flagellates by curvature inversion, like after light induced inversion of Flexa (Movies 1,2). Inhibition of Myo-II activity by ML-7 treatment led to the inhibition of hydrodynamically induced stimulation of inversion, showing a role of mechanically induced Myo-II activation at the origin of the hydrodynamically induced inversion (Fig. 5C,D,E and Movies 3,4). Strikingly, the inversion was additionally accompanied by an hydrodynamically stimulated Myo-II expression increase of a factor of 4 within the cell, 30 min after stimulation (Fig 5 F,G,H). This importantly demonstrates the existence of gene expression mechanical induction by the inversion in the pre-metazoa *flexa*, here having probably been selected to functionally strengthen and stabilize the inversion response.

This shows that wave-induced hydrodynamic mechanical stimulation triggers gastrulation-like inversion of the pre-metazoan multi-cellular choanoflagellate Flexa in an Myo-II-dependent manner.

## Environmental marine mechano-biochemical induction of first animal organisms emergence

Here, the present observations all together demonstrate that in addition to bilaterians (*11, 14-16*), mechanical stimulation of Myo-II dependent gastrulation is shared by cnidarians (*Nv*) and multi-cellular pre-metazoan choanoflagellates (Flexa). Given this lineage conservation, the lack of conserved biochemical signals involved in today’s metazoan early embryos Myo-II activation in gastrulation(*5-8*) and the fact that both *Nv* and Flexa gastrulation/inversion were hydrodynamically mechanostranductively induced by sea water wavelets, we propose that environmental mechanical strains such as marine flow and waves on the shore have been at the evolutionary origin of *urmetazoan* first animal organism emergence, due to the mechanosensitivity of Myo-II in *premetazoan* multi-cellular tissues more than 700 million years ago (Fig. 6a,b). This process could have been selected as a favourable reflex in response of feeding nutritive sediments that are found in suspension at necessarily higher concentration into the water in the presence of turbulent flows on the shore or on the sea ground, than without flow. A process even more favoured by the inherent transition to swimming, literally making multicellular choanofalgelatte “hunting swimming mouths” in response to flow induced favoured nutritive environment.

**Figure 6:**
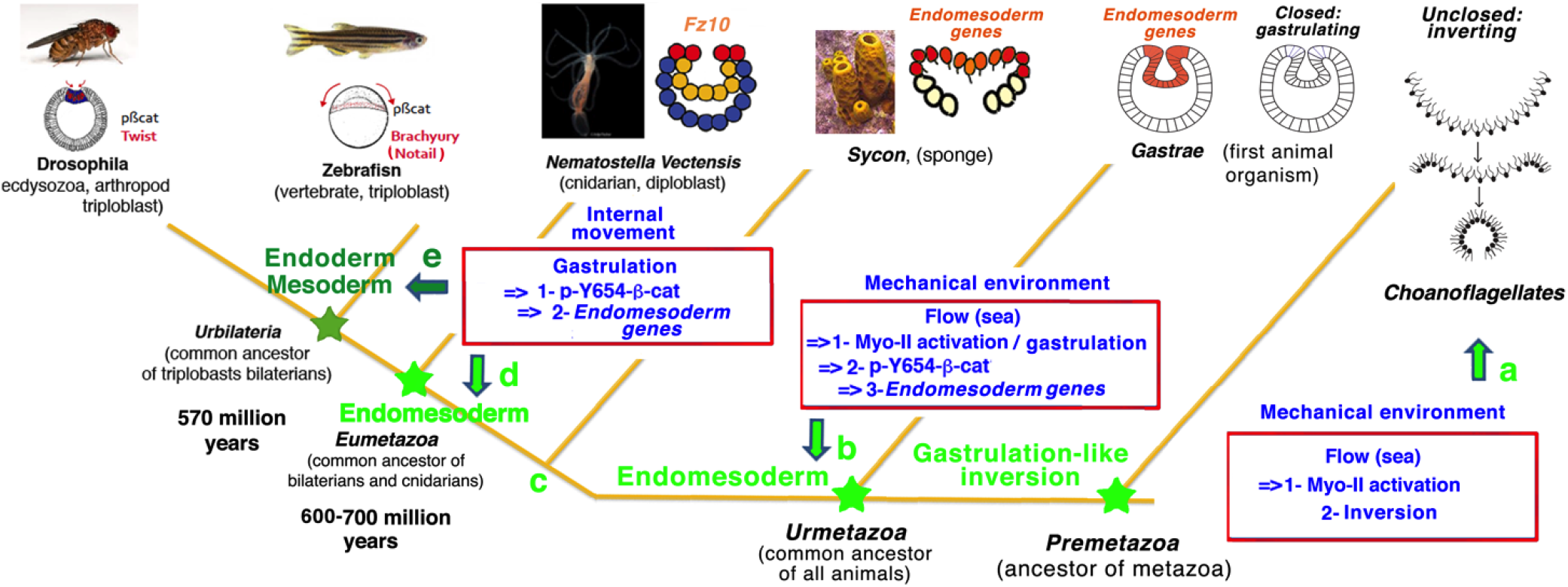
Mechanotransductive origins of first metazoan animal emergence evolutionary scenario. **a Results**-Pre-metazoa Flexa un-closed multi-cellular choanoflagelate inversion is mechanotranductively stimulated by hydrodynamic flow mimicking soft waves on the sea shore, in a Myo-II dependent mechanotransductive process (Fig.5), like *Nv* cnidarian gastrulation (Fig.1), and similarly to mechanotransductive activation of Drosophila bilaterian gastrulation(11). Scheme from (Brunet et al, 2019). **Evolutionary consequences**-Myo-II mechanotransductive induction of inversion/gastrulation is conserved all along metazoan evolution, from the hydrodynamically induced inversion of pre-metazoan unclosed multicellular choanofalgellates. **b Results**-Environmental hydrodynamic sea water hydrodynamic strains lead to mechanotransductive induction of gastrulation, with in turn gastrulation strains mechanotransductively stimulating endomesoderm *Fz10* gene expression in the gastrulating tissue of the Cnidarian *Nv*, in a Y654-β-cat phosphorylation activation mechanotransductive process conserved with bilaterians(9) (Fig. 3). **Evolutionary consequences**-Pre-metazoan inversion having led to hydrodynamic induction of gastrulation after closing of multicellular choanofalgellates, with gastrulation strains mechanotransductively leading to endomesoderm genes expression in gastrulating tissue after evolutionary emergence of β-cat mechanosensitivity, as conditions of the ur-metazoan first metazoan emergence. **c,d Results**-Internal morphogenetic movements of gastrulation lead to endomesodermal *Fz10* gene expression in the *Nv* cnidarian embryo in a Y654-β-cat phosphorylation dependent mechanotransductive process (Fig. 3). This is similar to the expression Twist and *brachuyry* endoderm and mesoderm genes in the early Drosophila and zebrafish bilaterian embryos(9). **Evolutionary consequences**-Genetically regulated internal morphogenetic movements (gastrulation) have replaced external environmental (*i.e.* sea water) mechanical strains in order to stimulate gastrulation and endomesoderm specification in the Eumetaoan common ancestor of cnidarian and bilaterian, thereby having initiated autonomous metazoan embryogenesis. **e**-The latter property has been conserved in both mesoderm and endoderm specification, after endoderm/mesoderm divergence at the origin of first bilaterian emergence, as demonstrated in early Drosophila and zebrafish embryos having directly diverged from the common ancestors of bilaterians(9).

From this point of view, the light-induced gastrulation of Flexa probably appeared later in the course of evolution and could have favoured swimming and feeding of choanoflagellate colonies at sea surface where light and plankton are both more abundant.

In contrast to open spheres (*Flexa*) closed spheres (*Nv*) cannot invert Our results further suggests that metazoans could have evolved from multicellular non patterned choanoflagellate to pre-patterned spheres with polarized high sensitivity of Myo-II to mechanical strains (patterned by Dsh in *Nv* (Fig. 2C,D), or Fog downstream of Dorsal in Drosophila (*27*)). Which would have allowed local inversion in a closed blastulae, instead of global inversion in an opened blastulae, namely gastrulation. We additionally propose that the emergence of genetically regulated internal morphogenetic movements have subsequently substituted environmental cues (such as *sna* dependent apex pulsation (*11*) in Drosophila, to be determined in *Nv*), thereby initiating autonomous embryogenesis independently of the environmental stimulation (Fig. 6c,d,e).

These observations additionally demonstrate that the mechanical stimulation of Y654-βcat phosphorylation in junctions that initiates EM genes expression by endogenous morphogenetic movements of gastrulation is shared by cnidarians (*Nv*) and bilaterians (zebrafish and *Drosophila*) (9) (Fig. 6d,e). Interestingly, *brachyury*, which was proposed to separate the endoderm from other tissue types including the ectoderm in *Nv* (*28*), has also been found as mechanosensitive in response to artificial non physiological deformation (uniaxial and 16 hours of application) applied to early *Nv* embryos in a β-catenin dependent way (*29*). However, due to the absence of junctional βcatenin-Cadherin complexes in premetazoan choanoflagellates (30) this mechanism should be absent in response in ancestral gastrulation-like inversion (Fig. 6a). Interestingly, the βcat-dependent activation of the EM *brachuyry* gene is activated during inversion in sponges embryos (Fig. 6c) (*31*). We thus suggest that the emergence of junctional βcat containing the major Y654 interactive site of βcat with Cadherins, and of adhesive Cadherins able to interact with βcat in *premetazoa* multicellular choanoflagelattes, might have initiated conditions for the emergence of mechanotransductively induced EM specification in response to environmental hydrodynamic mechanotransductive induced gastrulation or inversion, in the first metazoans (*urmetazoa*, Fig. 6b). These conditions might have initiated the transition from choanoflagelatte to sponges (32). Furthermore, this property, inherited by the 600-700 million years ago common ancestor of cnidarians and bilaterians, is still shared by metazoan early embryos at the onset of biomechanical morphogenesis (Fig. 6b-e), in contrast to the multitude of un-conserved other signalling processes that have subsequently added up in the course of evolution (*9, 23*) (Sup Info 8).

The presence of bacteria *per se* was proposed to have favoured first transient forms of small multicellular rosette aggregates inherently coupled to feeding functions (*33*). Here we propose that hydrodynamically induced gastrulation-like and motility stimulation behaviour of stabilized multicellular pre-metazoan, have favoured nutriment feeding. And that such primitive behaviourial process has initiated gastrulation, then EM specification, thereby led to early animal organisms evolutionary emergence. Both of which having subsequently been triggered by internal mechanical stimulation that have initiated autonomous animal embryogenesis during next steps of early animals evolution. With such lineage conservation from bilaterian to cnidarian and pre-metazoan importantly favouring the evolutionary origin hypothesis rather than a convergence evolution process.

## Author contributions

MN performed all hydrodynamical stimulation experiments on *Nv*, including hydrodynamic conditions setting-up, morphological phenotype analysis, *in situ* and IF labelling with and without drugs, and Dsh-DIX injections. MN and AJ realized embryo deformation measurement experiments. TM set up the *Nv* injections in the lab and performed stb-MO injection experiments, which consisted in the initiation of the present research. TM set-up imaging conditions of Y654-βcat and actin labelled embryos, and produced the algorithm for cell resolution image analysis with FS. EF performed the hydrodynamic experiments on Flexa. FB performed in vitro RNA transcription to produce antisense RNA probes used for in situ hybridization experiments on *Nv* and carried out Flexa labelling after fixation on glass substrate or in suspension. Y654 inhibitor experiments on Y654 phosphorylation and βcat IF labelling of non-hydrodynamically stimulated embryos were carried out by ACB. ACB set-up Y654-βcat labelling conditions and quantification on *Nv*. ER together with MN, performed part of the *Nv* injections and image analysis. JLG participated to the flow-meter experiments. EF coordinated the work, and wrote the first draft of the manuscript, implemented by all of the article authors.

## Acknowledgements

The authors thank Aldine Amiel, Hereroa Johnston, João Carvalho, Adrien Bouclet, Elena Fernandez-Sanchez, and Mark Martindale for training, scientific discussions, and biological material sharing, sending and setting up, David Quéré for scientific discussions and providing the rapid camera physical set-up, Thibaut Brunet for p-βcat labelling initiation tests in *Nv* and providing Flexa and culture protocols and discussions, Aude Batistella for help in Flexa culturing. This work was supported by the OCAV grant n° ANR-10-IDEX-0001-02 PSL, and EF lab is supported by the FRM (grant Équipe labellisée FRM 2015 DEQ20150331702, grant Équipe labellisée FRM 2019 EQU201903007805), the Inca (PLBIO-13-172 and PLBIO-03-ICR-1), the ANR (16-CE14-002801) and the LabEx Cell(n)Scale (grants ANR-11-LABX-0038, ANR-10-IDEX-0001-02).

## Data and materials availability

All data is available in the main text or the supplementary materials.

