## Supplemental Data for "Evolutionary Emergence of First Animal Organisms Triggered by Environmental Mechano-Biochemical Marine Stimulation"

**This PDF file includes:**

Materials and Methods
Supplementary Text
Figs. S1 to S4
Captions for Movies S1 to S4

**Other Supplementary Materials for this manuscript include the following:**

Movies S1 to S4
Code of cell scale image analysis

### Materials and Methods

#### *N. vectensis* culture and spawning

*Nematostella* sea anemones (WT, males and females, more than 1 year old, accession number ABAV01000000) are kept in artificial seawater (ASW) at 18°C in the dark and fed every day with fresh *Artemia* nauplii. Females and males are kept separately in different glass bowls (ϕ15x6cm) half-filled with ASW made by dissolving 10g of Instant Ocean salt (Aquarium Systems) in 1L of deionized water (1/3x SW). In each bowl are cultivated approximately 30-40 individuals.. Animals are moved to clean bowls with fresh 1/3x SW every two weeks. No further care is required during weekend.

Adult animals must be prepared for spawning every three weeks. From 24 to 48h before the day of spawning, animals are fed with defrost oyster. The night before spawning, animals are heated (~25°C) and lighted for 9 hours. After that they are moved to new bowls with fresh, cool (18°C) 1/3x SW. Sperm and eggs are collected separately. Oocytes are fertilized *in vitro* to synchronize development by dropping sperm into eggs 'beaker which is then gently shaken for 15 minutes at room temperature. The fertilized eggs are kept in dark at 18°C. In this condition, most embryos reach blastula stage after 18 hours post fertilization (hpf)

#### Hydrodynamic stimulation experiment

Liquid silicone based elastomer (Sylgard 184, Dow Corning) and curing agent was mixed at 10:1 ratio and then poured into ϕ 10mm Falcon petri dish (4g of mixture/dish). The dishes were left on a 37°C heater for 3h. When the mixture was almost dried out but remain sticky, sand with average grain size of 1mm was added and spread out to fully cover the dish floor. Sanded dishes were rinsed thoroughly with MilliQ water and then 1/3x filtered sea water (FSW) prior to experiment. These dishes can be reused.

At 16hpf, embryos were transferred to SPD filled with 22ml of 1/3x FSW. The dish was then put on shaker's plateaus (Major Science MS-NOR-30) which rotated at 85 rounds per minute and reversed the direction of rotation every 2.5s so that the movement mimicked the sea wavelets on shore for 2 h, from 16 to 18hpf. All experiments were carried out at 18°C unless otherwise stated.

Jellied embryos were then de-jellied using L-Cystein solution (0.4g/10ml of 1/3x FSW, pH adjusted to 7.4-7.6).

#### Rapid camera deformation imaging

The videos were recorded using a uEye camera equipped with a Navitar 4X Zoom lens, providing a depth of field on the order of 100 μm. The movies were shot at 10 frames per second with a resolution of 1 micrometer per pixel. A petri dish containing embryos was put on an LED light source which was attached to the shaker's plateaus. The whole system was placed under the camera. After one hour of stimulation, top-down 30-second-videos were captured every 5 minutes in order not to overheat samples. The images extracted from the videos showed how embryos deformed in response to the stimulation. This deformation is measured by their length-to-width ratio.

**ML-7 inhibitor**

ML-7 inhibitor (Sigma I2764) stock solution was prepared 50% (v/v) ethanol at concentration of 22mM. ML7 treatment was carried out at 14hpf and at final concentration of 10µM until the end of experiment.

**Phosho-Y654-βcat inhibitor**

In order to identify an Y654 βcat inhibitor, we tested 10 different compounds from Chem Div and first screen for toxicity and effective inhibiting Y654 phosphorylation. The compound n° S381-0393 gave the most consistent results at 100µM final concentration.

It is worth noting that this compound was dissolved in DMSO for reason of good solubility. Only for experiments followed by an *in situ* hybridization labelling, due to the fact that DMSO perturbed fz10 expression, the compound was exceptionally put in suspension at the same concentration in ASW instead.

**Rhodamin-Phalloïdin staining**

Embryos were fixed in fresh 4% paraformaldehyde (PFA) supplemented with 0.2% glutaraldehyde in 1/3x FSW for 30 min at room temperature. Fixed embryos were thoroughly rinsed in PBS-Tween 1% (PTw) and then incubated in Fetal Bovin Serum (FBS) 10% (v/v) . Finally, embryos were incubated in 300x diluted Rhodamin Phalloidin (Life Technologie, R415). Images were taken in black and white, and colored in green for a better visualisation in the Figures.

**β-catenin and p-β-catenin staining**

On the first day, fixed embryos were rinsed in PTw followed by FBS 10% (v/v) for 30 min. Then embryos were incubated in the primary antibody (anti-β-catenin produced in rabbit (Sigma, C2206) or anti-p-β-catenin produced in mouse (Clinisciences, NB-22-0209-S and 1:250, Santa Cruz Biotechnology (1B11) Ref 57533)) for one night at 4°C.

The next day, embryos were thoroughly rinsed in PTw and again in FBS 10% for 30min. After that, embryos were incubated in secondary Alexa Fluor 488 conjugated antibody. Rhodamine-Phalloidin was also added in this step together with the secondary antibody.

***In-situ* hybridization of *fz10* gene**

***Fixation***

To conserve at best the genome of embryos, we followed another protocol of fixation (Genikhovich&Technau DOI: 10.1101/pdb.prot5282). Briefly, embryos were rinsed inPTw, then rapidly pre-fixed in PFA 4%+ Glutaraldehyde 0.8% for 90s and finally fixed in PFA 4% for 2 h. Fixed embryos are then rinsed 3x in PTw, 2x in sterile water, 1x in pure Methanol. All steps are done on ice. Embryos are stored at -20°C in pure methanol for at least one night.

***Preparation of mRNA probe***

pCS2 DNA recombinant plasmids were used as templates to amplify the cDNA inserts by PCR. The PCR products were used as templates to synthesize anti-sense RNA probes containing digoxygenin-11-UTP (Roche Biochemicals).

*In vitro* transcription were carried out using the T7 Ambion message machine kit (Ambion, USA) to synthesize anti-sense probes for *Fz10, bmp-1 like* and *DIX-Dsh.* After DNAse RNAse-free treatment, RNA were dissolved in RNAse-free water.

***Procedure***

We followed the *in situ* hybridization protocol optimized by E. Rottinger's team(*17*).

**mRNA microinjection**

***Preparation of mRNA***

PCS2 Nv Dsh DIX GFP recombinant plasmid was linearized with Not1 restriction enzyme. The linearized template was used to synthesize capped mRNAs *in vitro* with the T7 Ambion message machine kit (Ambion, USA). Capped mRNAs were purified with the Ambion MegaClear kit (Ambion, USA), before starting injections.

mRNA diluted in RNAse free water at desired concentration was denatured by heating at 65°C for 10min. Then rhodamine-dextran fluorescent marker was added to injection solution at the concentration of 0.5µl/5ml. The control embryos were injected only with a mixture of sterile RNAse water and rhodamine-dextran.

Stb-MO was designed following reference (*7*).

***Microinjection experiment***

Embryos right after being fertilized and dejellied are maintained in 1/3x FSW at 16°C in order to slow down the development. Injection must be done within first 4 hours post fertilization, before the first cleavage. For injection experiment, embryos were deposited on a Falcon petri dish (351007) along some parallel scratches which served as landmarks when manipulation under microscope. These dishes are recommended for good adhesion of embryos on the floor. We used home-made micro-needles fabricated by a capillary pulling machine. The Femtojet microinjecter enables one to adjust a constant permanent pressure and injection pressure. In practice, these parameters are tuned so that the liquid volume at the needle end when injected makes a spot of diameter roughly 20% of embryo diameter.

Injected embryos are kept at 20°C. In this condition embryos reach normally blastula stage at 20hpf. Before any further experiments, we select only successfully injected embryos by removing all broken or without detectable fluorescent marker embryos.

Hydrodynamic stimulation was thus applied between 18 and 20 hpf embryos for Dsh-MO experiments.

### Imaging and analysis

#### *Spinning and microscope*

Fluorescent imaging and morphological observation was performed with Yokogawa spinning dish confocal (CSU-X1) coupled to Olympus inverted microscope (IX70). The actin, nucleus or β-catenin of Nv embryos shown in this article were either an image extracted from a stack or a MAX intensity projection of a whole stack obtained with ImageJ software.

Y654-β-cat phosphorylation quantitative analysis at the tissue scale were also performed by Image J. Images stacks (taken every two microns) of entire embryo which were summed. The total intensity/pixel of the whole embryo was measured by selecting the embryo only, and normalized to the average intensity of the external background. Intensity gradient was measured on the selection of the intensity's blastopore (invagination area), normalized by the intensity of the rest of the embryo. Background was subtracted for intensity measurement before normalisation. At 18 h of development, non-gastrulated embryos were orientated along the direction of maximum gradient.

Y654-β-cat phosphorylation quantitative analysis at the cell scale was performed by an algorithm using the phalloïdin junctional signal to discriminate between two different cells (see Code), after imaging stacks of images separated by 1 micron, of each embryo along its 3 principal axis (*34*).
To precisely analyse large numbers of embryos at the cell scale, we created an algorithm using Python programming. 3D images (stack) were projected to get 2D images (sum of 5 pixels around the z maximum). We first applied an Otsu filter to discriminate the background signal from the embryo signal. Then, based on the actin signal, we realized a skeletonization using the DisPerSE software (Sousbie, 2011). The skeleton was dilated up to 5 pixels to correspond to the width of the cell junctions. A watershed-based segmentation allowed us to create a cell catalogue reporting for each cell its apical area and its apical perimeter. Also for each channel (actin and pY654-bcat signals), the catalogue contains the mean intensities of the junctional and cytosolic cell signals respectively. All these measures were normalized to the background mean intensity.

Hybridization images were taken with a CoolSNAP (HQ2) camera, mounted on an upright widefield Leica microscope.

### *Choanoeca* Flexa

#### *Culture*

The Flexa strain was obtained from Thibaut Brunet (UC-Berkeley) and cultured in 10ml sea water medium in a sea salt 16.45g/l culture flask, in the presence of Rifamycine 20µg/ml antibiotic to inhibit bacterial dependent photosensitive inversion stimulation(*26*), with the difference that the antibiotic was directly diluted into the sea water medium without DMSO. Culture without the rice grain let to around 70 micron half-spheres, whereas as culture with the rice grain(*35*) let in addition to numerous 500 to 1000 microns tissues mixing half-spheres and almost complete spheres.

#### *Hydrodynamic stimulation*

Rice cultured Flexa were hydrodynamically stimulated after 4 to 5 days of culture, for 2min directly in there flasks, by the (Major Science MS-NOR-30) shaker used for *Nematostella* hydrodynamic stimulation, with a speed of 105 rpm and a positive to negative cycling period of 2.5s, leading to a flow on the order of 20 cm/s, after having removed the rice grain.

*ML7 treatment*

10 ml of the sea water cultured Flexa was implemented with 1 to 22.5μl of a 22mM concentration of ML7 suspended in 50% water and 50% ethanol, for 30min before hydrodynamic stimulation. Note that ML7 treatment did not perturb un-stimulated Flexa structures, but systematically weakened hydrodynamically stimulated Flexa structures, leading to smaller pieces of multi-cellular tissues after hydrodynamic stimulation.

*Morphological observation*

The Flexa morphological phenotypes were observed in transmission with a DMIRB inverted microscope Leica recoded with a C4742-95 Hamamatsu camera. Phenotype counting was initiated 20 minutes after the end of the stimulation.

*Fluorescent labelling and imaging.*

Flexa were fixed following reference procedures[21] as described in Brunet et al. 2019, on FluoroDishes and on mini glass slides pre-treated with a handheld Corona surface treater (Electro-Technic Products BD-20AC), coated with poly-D-lysine (Sigma Aldrich P6407-5MG). Imunofluorescence staining was performed using Rabbit anti-Myosin (1:10, M7648 Sigma Aldrich), and Rhodamin Phalloidin (1:100, Thermo Fisher Scientific) primary antibodies. Secondary antibodies : anti-rabbit Alexa 488 (1:300, Invitrogen) ; anti-mouse Alexa 647 (1:300, Invitrogen) were used. Sections were cover-slipped with Prolong Gold Antifade (Thermo Fisher Scientific). Images were taken with a Nikon A1R 25HD confocal microscope at the Nikon Imaging Centre located at the Curie Institute. Image were processed using Fiji/ImageJ software. Note that inverted structures, and ML7 treated structures revealed much more fragile than non inverted structures with regard to fixation, showing small patches of tissues only for inverted conditions, and no multi-cellular structure at all in ML7 treated conditions.

Note that inverted structures, and ML7 treated structures revealed much more fragile than non-inverted structures with regard to fixation, showing small patches of tissues only for inverted conditions, and no multi-cellular structure at all in ML7 treated conditions.

*Statistical test:*

Statistical tests used were either exact Fisher test (for number of positive and negative embryos counting) and the non-parametric Mann-Whitney test (for quantitative analysis associated to each individual embryo). In the specific case of Figure S1Ca in which a bimodal distribution with an important difference between the two populations appears in the un-jelled stimulated condition, the 5 elements of the upper population were compared to 5 elements representative of the mean value of the downer population. Sample size was empirically increased until an initial tendency in a first experiment could be confirmed by a p-value $<0.05$ in a replicate. All experiments where replicated at least 2 times (biological replicates).

Source Data: http://xfer.curie.fr/get/1sCB3jVEXqR/Source%20Data.zip

ICMJE guideline was followed.

**Supplementary Text**

**Sup Info:**

Sup Info 1:
Dejellied embryos were hydrodyamically stimulated following the same procedure than jellied embryos, except than sand was removed as individual dejellied embryos staid blocked against sand grains during stimulation. The deformation of the blastulae was monitored by fast imaging, and also found to be elliptic (Fig. S1Ca). Elliptic deformation was found to be of a 25-40% for 7% of the embryos at a given time (Fig. S1Ca-right-blue ring)
As seen in Fig. S1Cb-wavelets, we observed that gastrulation was initiated at 18h in nearly 30% of the stimulated embryos with a normalized *d/D* invagination depth equal or larger than 20% and apex constriction observed in the invaginating tissue (Fig. 1Cb, orange array), in contrast to static control in which 8% only of the embryos have initiated gastrulation.

The gel being of 1.5 to 4 mm radius, compared to the ~ 0.1mm radius of *Nv* embryos, hydrodynamic Stokes forces, proportional to the radius, should be on the order of magnitude 10 times more important on the gel embedding embryos compared to individual de-jellied embryos. Assuming an elastic modulus equivalent for the gel and for embryos, one expects a deformation of embryos significantly more important in jellied embryos, than in un-jellied embryos, in addition to potential sand friction effects on the ground. Which is observed in Fig.1C, and explains the higher rate of gastrulation response of jellied embryos experiments compared to dejellied embryos experiments.

Sup info 2:
Note that to our knowledge, no Myosin-II antibody was found to efficiently label *Nv* Myo-II thus preventing us to also follow the Myo-II behaviour in this specie.

Sup info 3:
Because injections can only be done in dejellied embryos, the role of Dsh in hydrodynamic stimulation of gastrulation initiation was tested on un-jelled embryos.

Sup Info 4:
i- Because the overall levels of pY654-βcat increases in the embryo, the oral-aboral gradient of pY654-βcat increase cannot be interpreted by an pY654-βcat decrease in the aboral domain of the embryo, and is due to an increase in pY654-βcat in the oral EM (Fig. S4A).
ii- Phosphorylation of Y654-βcat is classically low, and detectable after 3 hours of gastrulation only at the tissue scale in *Nv* (Fig. 4A), preventing us to check for Y654-βcat phosphorylation after the 2 hours of hydrodynamically stimulated gastrulation in 18hpf embryos protocol of all experiments.

Sup info 5:
Injecting stb-MO, that has previously been described to prevent *Nv* gastrulation (7), slightly decreased Y654-βcat phosphorylation total levels (as detected with cell scale imaging analysis) correlatively to the partial decrease in the number of invaginating and constricting embryos at 24h (the time at which stb-MO phenotypes most clearly distinguishes from WT phenotypes) (*34*), consistent with Y654-βcat phosphorylation sensitivity to gastrulation found in gastrulation defective ML7 treated embryos (Fig.S4B).

Sup Info 6:
Note that DMSO perturbed the expression of *fz10* in 18h embryos, such that Inhib-Y654-βcat was used at higher concentration without DMSO (see methods).

Sup info 7:
At gastrulation stage, β-cat nuclear staining was localised in single nuclear and peri-nuclear spots, co-localized with Dapi, as well as sometimes found as condensed out of the nucleus, as observed in other systems (Fig. 4 E-H) (*36*).

Sup Info 8: Mechanotransduction is involved in many physiological and pathological processes (*24, 37-46*). Into more details, we here propose that mechanotransductive cues have been re-enforced by biochemical patterning subsequently during the course of evolution, before the onset of morphogenetic movements (for instance by Dsh/Wnt in the *Nv* Cnidarian or by Dorsal in the Drosophila bilaterian arthropod) (*9, 23*) or after (for instance by Wnt, Bmp2 or Nodal found in the zebrafish bilaterian vertebrate) (*9*). Given the non-conservation of these biochemical signalling pathways in EM specification and invagination across metazoans (*5-8*), mechanical cues, which we show here to be evolutionary conserved from pre-metazoan choanoflagellates and cnidarians to bilaterians for EM gastrulation morphogenetic movements and specification respectively, have thus a high probability to be at the evolutionary origin of EM formation.

**Figure S1:** ***Nv*** **hydrodynamic stimulation conditions**

**A** Substrate of hydrodynamic stimulation with sand glued on pdms ground. **B** Embryos deformation in jelly. **C a** Deformation of un-jellied embryos, and quantification (Mann-Whitney statistical test). **b** Gastrulation of embryos submitted to hydrodynamic stimulation (N=2), and quantification based on significant normalized invagination depth *d/D* measurement higher or equal to 20% (see text, Fisher statistical test).

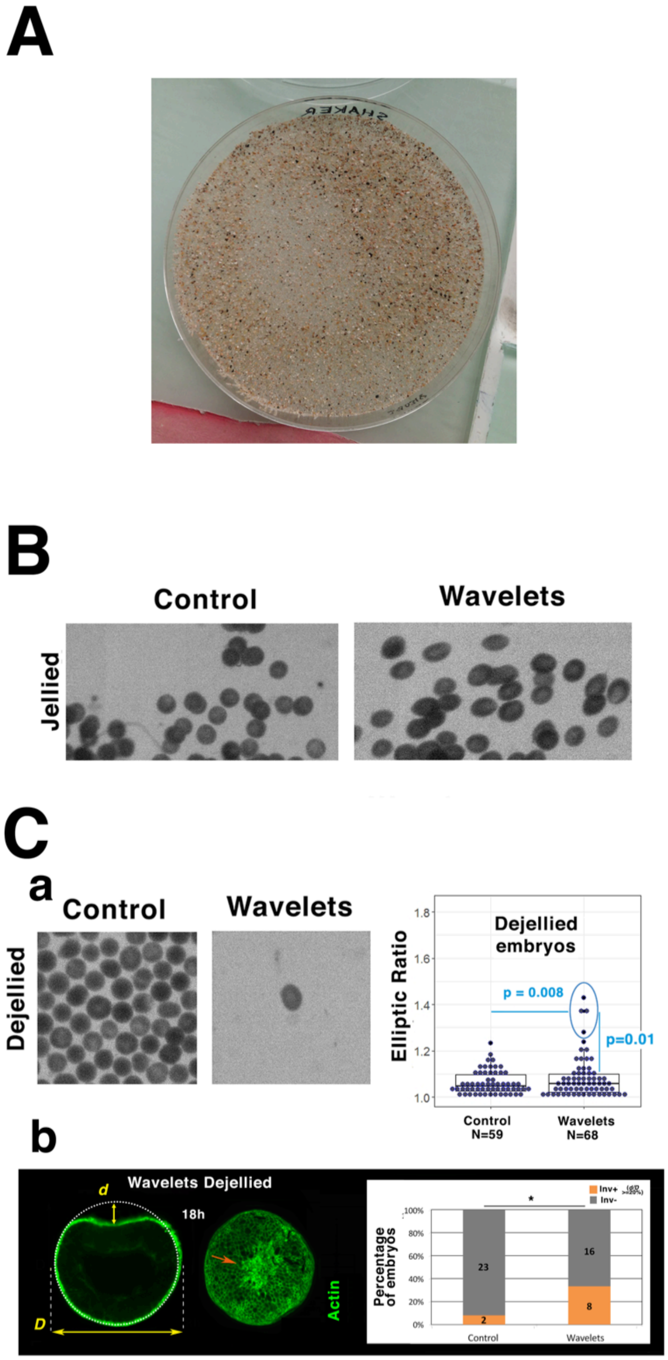

**Figure S2: bmp1-like is not stimulated by hydrodynamic treatment**
**A** 18hpf control and18hpf hydrodynamically stimulated. **B** Quantitative analysis with no increase, rather a decrease, of *bmp-1like* expression under hydrodynamic stimulation Statistical test: Fisher.

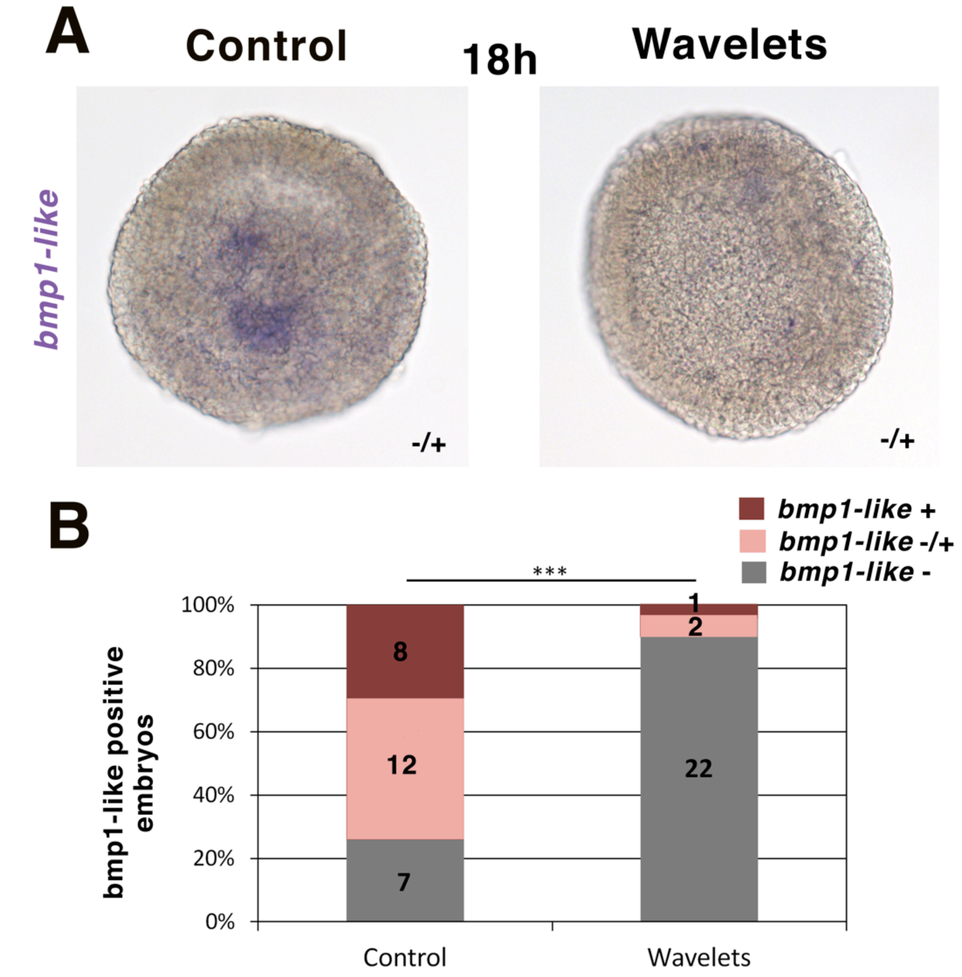

**Figure S3: Hydrodynamic stimulation of *Nematostella* embryos gastrulation by a flow-meter. A** Scheme of the flow-meter set-up. **B** Flow-meter induced gastrulation of 18hpf Nv embryos. **C** Quantification, Fisher statistical **test. N=2.**

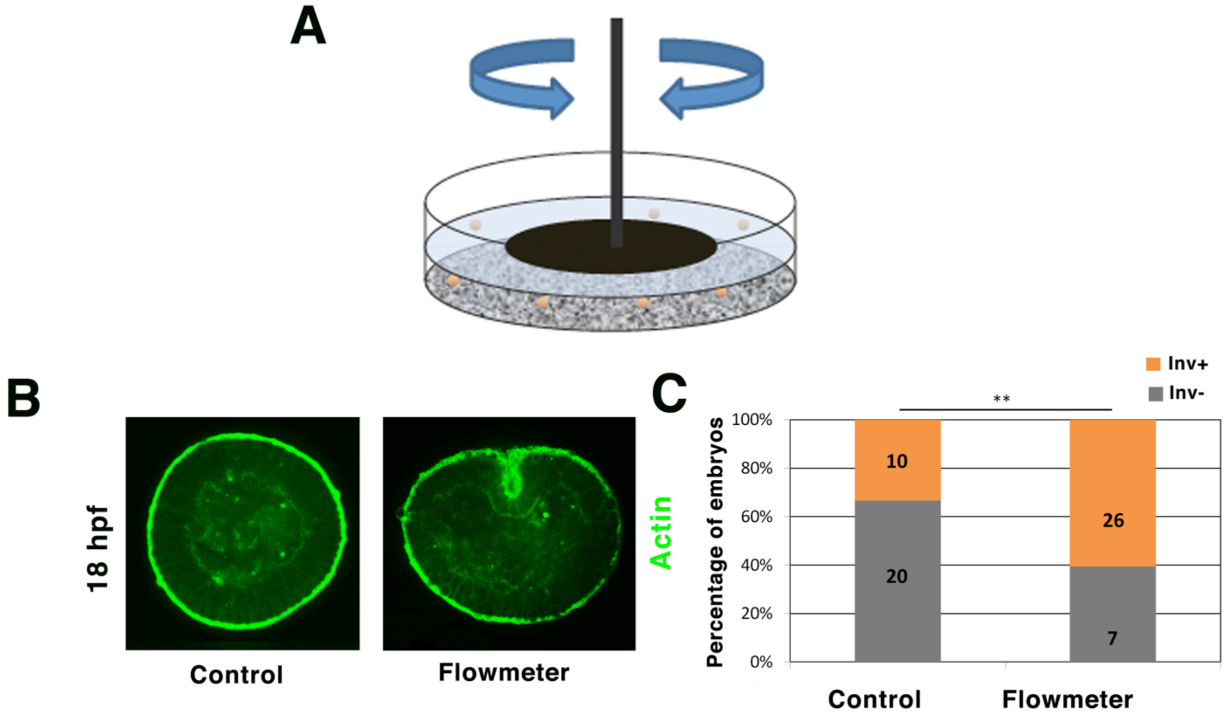

**Figure S4: Total pY654βcat levels quantification. A** 18h non-gastrulating and 21h gastrulating *Nv* embryos pharmacologically treated with p-Y654βcat inhibitor and ML7 Myo-II inhibitor**.** Values are normalized to the 18h control for total intensity. Statistical test: Mann-Whitney. N=2. **B.** 24h stb-MO injected embryos defective in gastrulation ($p<0.05$, exact Fisher test). Normalised pY654-βcat cytosolic intensity analised with computational programming (see Material&Methods). Statistical test for pY654-βcat intensity: Mann-Whitney $p<10^{-3}$ N=3.

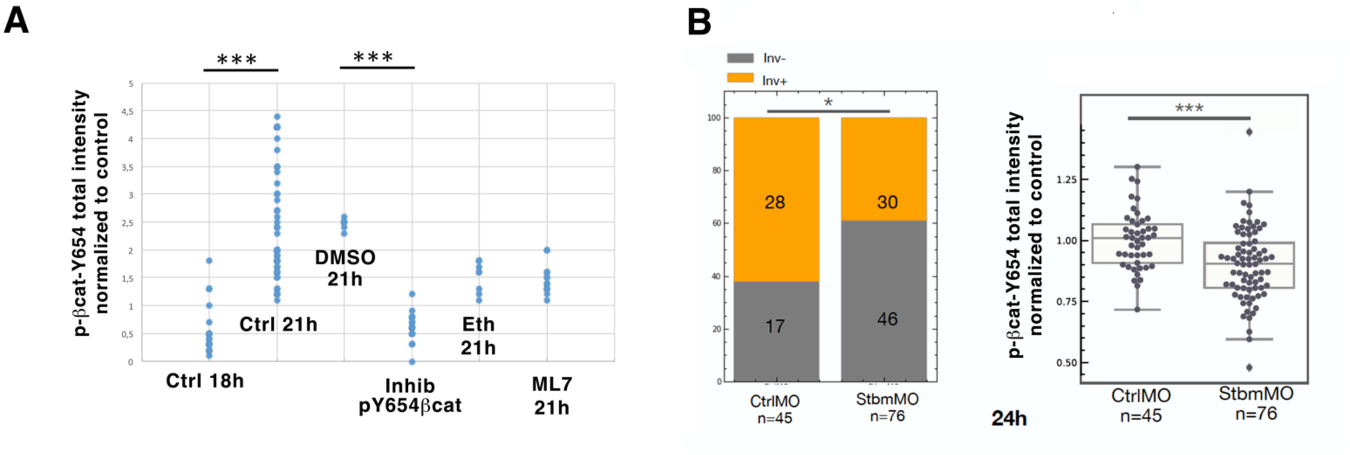

**Movie Captions:**

**Movie 1:** un-stimulated non-swimming spherical multi-cellular choanoflagellates, still opened at one pole. Changes of focal plan was induced in the movie to make sure that flagellates were not observed outside of the tissue.

**Movies 2:** stimulated swimming inverted multi-cellular choanoflagellates, **a** with a point of attachment to the substrate maintaining it in the same optical area **b** free and in this case larger and laterally opened. To follow it, the area of observation has to be changed several times.

**Movie 3:** stimulated swimming inverted multi-cellular choanoflagellates in the presence of the ethanol vehicle of ML7 alone, also probably punctually attached on the substrate.

**Movie 4:** stimulated non-swimming and non-inverted multi-cellular choanoflagellates in the presence of ML7. Changes of focal plan was induced in the movie to make sure that flagellates were not observed outside of the tissue.
