## Supplementary material for "Evolutionary Emergence of First Animal Organisms Triggered by Environmental Mechano-Biochemical Marine Stimulation": Code: Boxplot-Full.py

```
import numpy as np
import seaborn as sns
import matplotlib
import matplotlib.pyplot as plt
from scipy import stats
import scipy as sc
#import skimage
#from skimage import io
#from skimage.io.collection import ImageCollection,MultiImage
import glob
#from skimage import filters as filters
import os
#from skimage.morphology import disk
#from skimage.filters.rank import threshold
#from skimage.filters.rank import mean
#from skimage.filters.rank import enhance_contrast
import pandas
from joblib import Parallel, delayed
import multiprocessing
import pathlib


#VISUALISATION


enterin = sys.argv[1]
resin = sys.argv[2]

enter =  pandas.read_csv('../'+enterin, error_bad_lines=False)
label = [enter.columns.values[i] for i in range(len(enter.columns.values)) if i % 2 == 0]
levels = [enter.columns.values[i] for i in range(len(enter.columns.values)) if i % 2 != 0]
data = pandas.read_csv('../results/'+resin, error_bad_lines=False)
#TOTAL !

NAME = np.array(data['Name'])


act_all = np.array(data['Actine_all'])
act_fil = np.array(data['Actine_fil'])
act_outfil = np.array(data['Actine_outFil'])
bcat_all = np.array(data['Bcat_all'])
bcat_fil = np.array(data['Bcat_fil'])
bcat_outfil = np.array(data['Bcat_outFil'])

DATAT = []
DATAT.append(act_all)
DATAT.append(act_fil)
DATAT.append(act_outfil)
DATAT.append(bcat_all)
DATAT.append(bcat_fil)
DATAT.append(bcat_outfil)


for j, k in enumerate(NAME):
    for z, i in enumerate(NAME):
            if k[0:-9] == i[0:-9]:
                if DATAT[2][j] > DATAT[2][z]:
                    DATAT[0][z] = -0.21
                    DATAT[1][z] = -0.21
                    DATAT[2][z] = -0.21
                    DATAT[3][z] = -0.21
                    DATAT[4][z] = -0.21
                    DATAT[5][z] = -0.21


for plot in range(6):
    matplotlib.rcParams.update({'font.size': 6})
    fig = matplotlib.pyplot.figure(figsize=(6,7),dpi=100, facecolor='white')
    fig.subplots_adjust(wspace=.1,left=.01,bottom=.01)
    ax = fig.add_subplot(1,1,1,axisbg='w')
    matplotlib.rc('axes', linewidth=1)

    fontsize = 8
    
    for tick in ax.xaxis.get_major_ticks():
        tick.label1.set_fontsize(fontsize)
    for tick in ax.yaxis.get_major_ticks():
        tick.label1.set_fontsize(fontsize)

    
    DATA = DATAT[plot]

    X = []
    
    for i in label:
        fantome = []
        for j in enter[i]:
            for l,k in enumerate(NAME):
                try:
                    if j == k and DATA[l] != -0.21 and not(np.isnan(DATA[l])):
                        fantome.append(DATA[l])
                except TypeError:
                    if 1 == 2:
                        print('')                     
        X.append(fantome)
      
    mean = []
    median = []
    sd = []
    for i, j in enumerate(X):
            mean.append(np.mean(j))
            median.append(np.median(j))
            sd.append(np.std(j))
      
    dat = {'Name': label,
              'Moyenne': mean,
              'Median': median,
              'Sd': sd
}
    save = pandas.DataFrame(dat, columns=['Name', 'Moyenne', 'Median', 'Sd'])
      
    if plot == 0:
          save.to_csv('act_all.csv', sep=',')
    if plot == 1:
          save.to_csv('act_fil.csv', sep=',')
    if plot == 2:
        save.to_csv('act_outfil.csv', sep=',')
    if plot == 3:
        save.to_csv('bcat_all.csv', sep=',')
    if plot == 4:
        save.to_csv('bcat_fil.csv', sep=',')
    if plot == 5:
        save.to_csv('bcat_outfil.csv', sep=',')

    ref = X[0]
    DATA = []
    ax.set_ylabel('Normalized intensity', fontsize=10)
    for i in range(len(X)):
        X[i] = X[i]/np.mean(ref)
        for z in X[i]:
            DATA.append(z)

    #cet Y est labscisse (oui oui)   
    Y = [] 
    for i ,j in enumerate(label):
        p=0
        while j[p] != '_':
            p=p+1 
        k=0
        while j[k] != ' ':
            k=k+1   
        Y.append(str(j[0:p] + '\n' + j[p+1:k] + '\n' + j[k:len(j)] + '\n(n =' + str(len(X[i])) + ')'))
    
    #regarder sur internte toutes les options du bloxplot
    
    
    abscice=list(range(len(label)))
    meanX=np.zeros(len(label))
    sigX=np.zeros(len(label))
    SEMX=np.zeros(len(label))
    for i in range(len(label)):
        abscice[i]=abscice[i]
        meanX[i]=np.mean(X[i])
        sigX[i]=np.std(X[i])
        SEMX[i]=stats.sem(X[i])
        

    #plt.xlim(-1,len(label))
    plt.ylim(0.,np.max(DATA)+0.05+1*0.05*len(X)) #lim du plot en y. Mais attention, car lie au l220 a 239

#####On crée un dataframe pandas à partir de X (normalized intensity)

#X2 : liste avec le type de condition (+,- ou n)
#Xlabels : liste avec le nom de la condition globale (e.g. CtrlMO_24h)
    X2 = []
    Xlabels = []
    for i,l in enumerate(label):
         cond = np.empty_like(X[i],dtype=np.str)
         cond[:] = l[-1]
         X2.append(cond)
         if l[-1] != 'n':
             lab=l[:-5]
         else:
             lab=l[:-13]
         Xlabels.append([lab for j in range(len(X[i]))])
#On crée le dataframe pandas "df" à partir de X,X2 et Xlabels
    newdata = np.c_[np.concatenate(X),np.concatenate(Xlabels),np.concatenate(X2)]
    names = ['Normalized Intensity','condition','type']
    df = pandas.DataFrame.from_records(newdata,columns=names)
    df['Normalized Intensity'] = pandas.to_numeric(df['Normalized Intensity'])

#On merge les type de condition pour chaque condition globale (e.g. Inv+,Inv- et Constriction pour CtrlMO_24h)
    Xmerge = [np.concatenate(X[i:i+3]) for i in range(0,len(label),3)]

######SEABORN PLOT############
    import seaborn as sns
#sns.violinplot(data=X,bw=0.3, scale="count", inner="stick")
    green1 = ['limegreen']
    red1 = ['#BF0000']
    pal=sns.color_palette(red1)
    sns.boxplot(x='condition',y='Normalized Intensity',data=df,palette=pal)
    #sns.swarmplot(data=X,color=".25")
    colors3 = ['#030aa7','orange','#f4320c']
    pal2 = sns.color_palette(colors3)
    sns.swarmplot(x='condition',y='Normalized Intensity',data=df,hue='type',palette=pal2)
###############################
    #plt.xticks(abscice,Y)
    #for xtick, color in zip(ax.get_xticklabels(),pal):
    #    xtick.set_color(color)
    symbol = 'r*'

# ETOILE SIGNIFICANCE
    for i, j in enumerate(Xmerge):
        for k, m in enumerate(Xmerge):
            if i < k:
                a = sc.stats.mannwhitneyu(j,m)
                if a[1] < 0.0001:
                    plt.plot(k, np.max(DATA)+0.05+1*0.05*i , symbol)
                    plt.plot(k+0.1, np.max(DATA)+0.05+1*0.05*i , symbol)
                    plt.plot(k+0.2, np.max(DATA)+0.05+1*0.05*i , symbol)
                    plt.plot(k+0.3, np.max(DATA)+0.05+1*0.05*i , symbol)
                elif a[1] < 0.001:
                    plt.plot(k, np.max(DATA)+0.05+1*0.05*i , symbol)
                    plt.plot(k+0.1, np.max(DATA)+0.05+1*0.05*i , symbol)
                    plt.plot(k+0.2, np.max(DATA)+0.05+1*0.05*i , symbol)
                elif a[1] < 0.01:
                    plt.plot(k, np.max(DATA)+0.05+1*0.05*i , symbol)
                    plt.plot(k+0.1, np.max(DATA)+0.05+1*0.05*i , symbol)
                elif a[1] < 0.05:
                    plt.plot(k, np.max(DATA)+0.05+1*0.05*i , symbol)
                else:
                    plt.text(k, np.max(DATA)+0.05+1*0.05*i, 'NS', color = 'red', horizontalalignment='center')
    if plot == 0:
        plt.savefig('actine_all_swarn.svg', dpi = 600,  format='svg',bbox_inches='tight')
    if plot == 1:
        plt.savefig('actine_Fil_swarn.svg', dpi = 600,  format='svg',bbox_inches='tight')
    if plot == 2:
        plt.savefig('actine_outFil_swarn.svg', dpi = 600,  format='svg',bbox_inches='tight')
    if plot == 3:
        plt.savefig('bcat_all_swarn.svg', dpi = 600,  format='svg',bbox_inches='tight')
    if plot == 4:
        plt.savefig('bcat_Fil_swarn.svg', dpi = 600,  format='svg',bbox_inches='tight')
    if plot == 5:
        plt.savefig('bcat_outFil_swarn.svg', dpi = 600,  format='svg',bbox_inches='tight')

#afficher le graph apres analyse (ca prend du temps, ca sert a rien)
#plt.show()
```
