## Supplementary material for "Evolutionary Emergence of First Animal Organisms Triggered by Environmental Mechano-Biochemical Marine Stimulation": Code: MainAnalysis.py

```
# coding: utf-8
import os
import sys
import numpy as np
import scipy as sc
import matplotlib.pyplot as plt
from astropy.io import fits
from astropy.convolution import convolve,Tophat2DKernel
from PIL import Image
from pathlib import Path
import pandas
from termcolor import colored, cprint

import glob
from joblib import Parallel, delayed
import multiprocessing
import pathlib


import skimage
from skimage import filters as filters
from skimage.morphology import disk
from skimage.filters.rank import threshold
from skimage.filters.rank import mean
from skimage.filters.rank import enhance_contrast
from skimage.io.collection import ImageCollection,MultiImage
from matplotlib.colors import LogNorm
from scipy.ndimage.filters import gaussian_filter

### func def
def otsufilter(img):
    '''This function applies an Otsu filter, it takes in arguments a matrix and return a mask where the background is set to 0 and the foreground to 1. Change nbins with the type of images (8bits = 256, 16bits = 65536'''
    val = filters.threshold_otsu(img, nbins = 65536)
    mask = img < val
    mask = np.invert(mask)
    mask = sc.ndimage.binary_fill_holes(mask)
    return mask

def stack_importation(chemin):
    '''This function imports a stack of images .TIF in a 3D matrix, it takes in arguments the path of the folder.'''
    img = MultiImage(chemin)
    size = len(img)
    if (size != 1) :
        shape = img[0].shape
        stack = np.zeros((shape[0],shape[1],size))
        for i in range(size):
            stack[:,:,i] = img[i]
    if (size == 1):
        shape = img[0].shape
        stack = np.zeros((shape[1],shape[2],shape[0]))
        for i in range(shape[0]):
            stack[:,:,i] = img[0][i,:,:]
        shape=stack[:,:,0].shape
    
    return stack, size, shape
    

def number_MIP(matrix,shape,number):
    '''This function makes a N-maximums intensity projection of a 3D matrix, it takes in arguments the 3D matrix, the shape of the 3D matri in a tuple and the number of maximas to project.'''
    projection = np.zeros(shape)
    for x in range(shape[0]):
        for y in range(shape[1]):
            array_sort = np.sort(matrix[x, y, :])
            projection[x, y] = np.sum(array_sort[-number::])
    return projection


def proj_aroundMAX(matrix,shape,number):
    '''This function makes a maximum intensity projection of a 3D matrix, it takes in arguments the 3D matrix, the shape of the 3D matri in a tuple and the number of pints around the max to project.'''
    projection = np.zeros(shape)
    for x in range(shape[0]):
        for y in range(shape[1]):
            ind_max = np.argmax(matrix[x, y, :])
            inf = max(ind_max-number,0)
            sup = min(ind_max+(number+1),matrix.shape[2])
            projection[x, y] = np.mean(matrix[x, y, inf:sup])
    return projection

#Function to convert .tif to .fits
def tif_to_fits(base,chemin):
    im3d,im3d_size,im3d_shape = stack_importation(chemin)
    im = proj_aroundMAX(im3d,im3d_shape,2)
    hdu = fits.PrimaryHDU(im)
    hdu.writeto(base)
    return

#Main function
def MainAnalysis(enterin,resout,Workdir,imMaindir,MSCdir,KernelSize):
    if not os.path.exists(Workdir+'results'):
        os.makedirs(Workdir+'results')
    resultFile=Workdir+'results/'+resout

    enter =  pandas.read_csv(Workdir+enterin, error_bad_lines=False)
    data = enter.fillna(value=0)
    label = [enter.columns.values[i] for i in range(len(enter.columns.values)) if i % 2 == 0]
    levels = [enter.columns.values[i] for i in range(len(enter.columns.values)) if i % 2 != 0]


    cheminFace =[]
    glo_act=[]
    glo_act_fil=[]
    glo_bcat_out=[]
    glo_bcat=[]
    glo_bcat_fil=[]
    glo_act_out=[]
    notFOUND=[]
    
#LOOP ON CONDITION
    ii = 0
    for i, k in enumerate(label):
    #LOOP ON IMAGES
        for j, base in enumerate(data[label[i]]):
            if type(base) == str:
            
                print('%s' % (''))
                cprint(colored(base,'blue',attrs=['bold']))

                command = 'find '+imMaindir+' -iname "'+base+'.tif"'
                chemin=os.popen(command).read()[0:-1]
            #cprint(colored('   Dans le Dossier : '+chemin[0:-(len(base)+4)],'magenta'))
            #print('%s' % (''))
                fileIm_TIF=Path(chemin)
                if fileIm_TIF.is_file():

                ### 1. APPLY OTSU FILTER TO EMBRYO IMAGE

                    fileIm0=Path(Workdir+base+'_5max.fits')
                #fileIm=Path(base+'_Otsu.fits')
                    
                    if not(fileIm0.is_file()):
                        tif_to_fits(Workdir+fileIm0.name,chemin)
                    
                    original = fits.open(fileIm0)
                
                    maskOtsu=otsufilter(original[0].data)
                    originalIm=original[0].data
                    OtsuIm= maskOtsu*originalIm
                
                #if not(fileIm.is_file()):
                #    original[0].data = maskOtsu*original[0].data
                #    original.writeto(base+'_Otsu.fits')
                        
                ### 2. LAUNCH DISPERSE
                    level =str(np.int32(data[levels[i]][j]))
                    fileFil=Path(Workdir+fileIm0.name+'_c'+level+'.up.NDskl.fits')
                    fileMSC=Path(MSCdir+fileIm0.name+'.MSC')
                    if not(fileFil.is_file()):
                        if not(fileMSC.is_file()):
                        #os.system("source RunDisperse.sh "+base+' '+level)
                        #os.system("source RunDisperse-loadMSC.sh "+base+' '+level+' '+MSCdir)
                            os.system('mse '+Workdir+fileIm0.name+' -outDir '+Workdir+' -cut '+level+' -periodicity 0 -upSkl')
                        else:
                            os.system('mse '+Workdir+fileIm0.name+' -outDir '+Workdir+' -loadMSC '+MSCdir+fileMSC.name+' -cut '+level+' -periodicity 0 -upSkl')
                    
                        os.system('skelconv '+Workdir+fileIm0.name+'_c'+level+'.up.NDskl -outDir '+Workdir+' -toFITS')
                                                
                        
                     
                ### 3. ANALYSIS OF DISPERSE SKELETON
                #phBcat = fits.open('MAX_Nv-StbmMO+compression-20C-24hpf-E05-02_w2491.fits')
                
                    filaments = fits.open(fileFil)

                    filaments[0].data=maskOtsu*filaments[0].data
                
                #Put filaments value to 1 (0 otherwise)
                    filaments[0].data[np.where(filaments[0].data != 0)]=1.
                    
                #Convolve filaments with kernel to enlarge them
                    ConvFil = convolve(filaments[0].data,Tophat2DKernel(KernelSize))
                    ConvFil[np.where(ConvFil != 0.)] = 1.
                    
                #Apply filament mask to Otsu image
                    ImFil=maskOtsu*originalIm
                    ImFil[np.where(ConvFil == 0.)] = 0.
                    
                #Compute mean level of actine in filaments
                    Mean_ImFil=np.mean(ImFil[np.where(ImFil != 0)])
                    
                # Background using antiMaskOtsu
                    antiMaskOtsu = np.invert(maskOtsu)
                    BackgroundIm =  antiMaskOtsu*originalIm
                    MeanBackground_act = np.mean(BackgroundIm[np.where(BackgroundIm != 0)])
                
                
                # MeanFil_actine / MeanBackground
                    FinalMean_act_fil = Mean_ImFil/MeanBackground_act
                
#### ACTINE GLOBAL
                #Compute mean level of actin
                    Mean_Otsu_act=np.mean(OtsuIm[np.where(OtsuIm != 0)])
                #MeanAct/MeanBackground
                    FinalMean_act = Mean_Otsu_act/MeanBackground_act
                    
##### ANTI MASK TO ACTINE!
                    antiMask = np.zeros(ConvFil.shape)
                    antiMask[np.where(ConvFil != 0.)] = 2.
                    
                    antiMask[np.where(ConvFil == 0.)] = 1.
                    antiMask[np.where(antiMask == 2.)] = 0.
                    antiMask = maskOtsu*antiMask
                    
                    Im_antiMask=maskOtsu*originalIm
                    Im_antiMask[np.where(antiMask == 0.)] = 0. #ACTINE OUTSIDE FIL
                    
                #Compute mean level of actine outside filaments
                    Mean_act_out=np.mean(Im_antiMask[np.where(Im_antiMask != 0)])
                    
                # MeanFil / MeanBackground
                    FinalMean_act_out = Mean_act_out/MeanBackground_act
                    
### APPLY ANTI MASK TO B-CAT  
                    command = 'find '+imMaindir+' -iname "'+base[0:-5]+'w2491.tif"'
                    chemin=os.popen(command).read()[0:-1]
                #cprint(colored('   Dans le Dossier : '+chemin[0:-(len(base)+4)],'magenta'))
                #print('%s' % (''))
                    fileIm_TIF=Path(chemin)
                    if fileIm_TIF.is_file():
                        fileIm_b=Path(Workdir+base[0:-5]+'w2491_5max.fits')
                    if not(fileIm_b.is_file()):
                        tif_to_fits(Workdir+fileIm_b.name,chemin)
                    bcat = fits.open(fileIm_b)
                    bcatIm=bcat[0].data
                    maskOtsu_bcat=otsufilter(bcatIm)
                    Otsu_bcatIm = maskOtsu_bcat * bcatIm
                    
                    anti_bcatIm = antiMask * Otsu_bcatIm  
                    
                      #Compute mean level of bcat outside filaments
                    Mean_anti_bcatIm=np.mean(anti_bcatIm[np.where(anti_bcatIm != 0)])
                    
                      #Compute mean level of bcat
                    Mean_bcatIm=np.mean(Otsu_bcatIm[np.where(Otsu_bcatIm != 0)])
                      
                      # Background using antiMaskOtsu
                    
                    antiMaskOtsu_bcat = np.invert(maskOtsu_bcat)
                    Background_bcat =  antiMaskOtsu_bcat*bcatIm
                    MeanBackground_bcat = np.mean(Background_bcat[np.where(Background_bcat != 0)])
                      
                      
                      # MeanF / MeanBackground
                    FinalMean_bcat_out = Mean_anti_bcatIm/MeanBackground_bcat
                    FinalMean_bcat_all = Mean_bcatIm/MeanBackground_bcat
            
### APPLY FIL MASK TO B-CAT
                    fil_bcatIm = maskOtsu_bcat * bcatIm
                    fil_bcatIm[np.where(ConvFil == 0.)] = 0.
                    
                      #Compute mean level of bcat in filaments
                    Mean_fil_bcatIm=np.mean(fil_bcatIm[np.where(fil_bcatIm != 0)])
                    FinalMean_bcat_fil = Mean_fil_bcatIm/MeanBackground_bcat
                          
                          
                    cheminFace.append(base)
                    glo_act.append(FinalMean_act)
                    glo_act_fil.append(FinalMean_act_fil)
                    glo_act_out.append(FinalMean_act_out)
                    glo_bcat_out.append(FinalMean_bcat_out)
                    glo_bcat.append(FinalMean_bcat_all)
                    glo_bcat_fil.append(FinalMean_bcat_fil)
                #### APPEND ICI + CREE LQ LISTE VIDE L.74
                #grad.append(j[0])

                #delete temporary files (image.fits and disperse files)
                #if fileIm0.is_file():
                #os.system('rm '+fileIm0.name)
                    fileSkl=Path(Workdir+fileIm0.name+'_c'+level+'.up.NDskl')
                    if fileSkl.is_file():
                        os.system('rm '+Workdir+fileSkl.name)
                    fileMSC0=Path(Workdir+fileIm0.name+'.MSC')
                    if fileMSC0.is_file():
                        os.system('mv '+Workdir+fileMSC0.name+' '+MSCdir)

                    dati = {'Name': base,
                       'Actine_all': FinalMean_act,
                       'Actine_fil': FinalMean_act_fil,
                       'Actine_outFil' : FinalMean_act_out,
                       'Bcat_all': FinalMean_bcat_all,
                       'Bcat_fil': FinalMean_bcat_fil,
                       'Bcat_outFil': FinalMean_bcat_out}

                    #Le fichier results est reecrit a chaque fois quon refait tourner le programme. Si on veut le conserver, bien penser a le renommer.                      
                    save = pandas.DataFrame(dati, columns=['Name','Actine_all','Actine_fil','Actine_outFil','Bcat_all','Bcat_fil','Bcat_outFil'],index=[ii])
                    ii = ii + 1
                    if i == 0 and j == 0:
                        save.to_csv(resultFile, sep=',',header=True)
                    else:
                        save.to_csv(resultFile, sep=',',mode='a', header=False)
                else:
                    notFOUND.append(base)

#'Nomcol': nomliste,
#                dati = {'Name': base,
#                       'Actine_all': FinalMean_act,
#                       'Actine_fil': FinalMean_act_fil,
#                       'Actine_outFil' : FinalMean_act_out,
#                       'Bcat_all': FinalMean_bcat_all,
#                       'Bcat_fil': FinalMean_bcat_fil,
#                       'Bcat_outFil': FinalMean_bcat_out}


#Le fichier results est reecrit a chaque fois quon refait tourner le programme. Si on veut le conserver, bien penser a le renommer. 
#                save = pandas.DataFrame(dati, columns=['Name','Actine_all','Actine_fil','Actine_outFil','Bcat_all','Bcat_fil','Bcat_outFil'])
#                save.to_csv(resultFile, sep=',',mode='a', header=False)


#PRINT un résumé de ce qui a fonctionné ou non dans le terminal
    print('%s' % (''))
    print('%s' % (''))
    print('%s' % (''))
    cprint(colored(('%s' % ('###########################################')),'red'))
    cprint(colored(('%s' % ('                  RÉSUMÉ')),'red',attrs=['bold']))

    if len(notFOUND) != 0:
        print('%s' % (''))
        cprint(colored(('%s' % ('        !!ATTENTION !!')),'red',attrs=['bold','blink']))
        cprint(colored(('%s' % ('Les images suivantes n\'ont pas été trouvées')),'red',attrs=['bold']))
        print('%s' % (''))
        for name in notFOUND:
            cprint(colored(('%s' % (name)),'white'))
            print('%s' % (''))
            cprint(colored(('%s' % ('Verifiez bien qu\'elles sont dans le bon dossier ! ')),'red',attrs=['bold']))
            print('%s' % (''))
            cprint(colored(('%s' % ('!! Les *AUTRES* images du enter.csv ont été analysées sans erreur ! (a priori) !!')),'green'))
    else:
        print('%s' % (''))
        cprint(colored(('%s' % ('!! BRAVO, *TOUTES* les images du enter.csv ont été analysées sans erreur ! (a priori) !!')),'green',attrs=['bold']))
    cprint(colored(('%s' % ('Les résulats sont dans le fichier '+resultFile)),'white',attrs=['bold']))
    cprint(colored(('%s' % ('###########################################')),'red'))
    print('%s' % (''))
    print('%s' % (''))

    return

#Read arguments
if __name__ == "__main__":
    import sys
    MainAnalysis(sys.argv[1],sys.argv[2],sys.argv[3],sys.argv[4],sys.argv[5],float(sys.argv[6]))
```
