## Supplementary material for "Evolutionary Emergence of First Animal Organisms Triggered by Environmental Mechano-Biochemical Marine Stimulation": Code: README.txt

---- Intro and quick program description

The program MainAnalysis.py allows one to launch the full analysis of the embryo imuno-fluorescence signals from images taken in the BCAT wavelenghts (e.g. 491 nm) AND in the ACTINE wavelenghts (e.g. 561 nm).


Brief description of the code (from Nguyen, Merle et al. submitted):
To precisely analyse large numbers of embryos at the cell scale, we created an algorithm using Python programming. 3D images (stack) were projected to get 2D images (sum of 5 pixels around the z maximum). We first applied an Otsu filter to discriminate the background signal from the embryo signal. Then, based on the actin signal, we realized a skeletonization using the DisPerSE software (Sousbie, 2011﻿). The skeleton was dilated up to 5 pixels to correspond to the width of the cell junctions. A watershed-based segmentation allowed us to create a cell catalogue reporting for each cell its apical area and its apical perimeter. Also for each channel (actin and pY654-bcat signals) the catalogue contains the mean intensities of the junctional and cytosolic cell signals respectively. All these measures were normalized to the background mean intensity.


--------- Step 0. Prepare the input file

After having taken the images, name them as follow : EmbryoName_w3561.tif and EmbryoName_w2491.tif and put them somewhere in your image directory (which may contains several sub-directories for different experiments for example).

Before launching the program you need to create an input csv file to feed the program (that we will now call 'enter.csv').

An example of enter.csv is given in the test-data/ sub-directory.
There are 2 columns for each observed phenotype :
1. the first column contains the name of the embryos (with the '_w3561'extension, without '.tif')
2. the second column (no name) contains the detection level that Disperse will use.


You can do as many pairs of columns as you want in a single enter.csv file. The different pairs of columns do not need to have the same number of lines. These pairs of columns can be used to analyse separately different observing conditions (e.g. [control group vs test group] OR [phenotype1 vs phenotype2 vs phenotype3]).


------- Step 1. Launch the main program

To launch the program, use the following syntax :

python3 MainAnalysis.py arg1 arg2 arg3 arg4 arg5 arg6

arg1: name of the input csv file e.g. 'enter.csv'

arg2: name of the output csv file e.g. 'results.csv'

arg3: working directory i.e. where the 'enter.csv' file is located and where your results will be saved

arg4: root directory where the program will look for the images. Images can be located in sub-directories inside this root directory

arg5: directory where the .MSC file (saved and the re-used by disperse) are (or will be) located

arg6: Size of the kernel to be applied to enlarge filaments in number of pixel. Recommended value is 2 (corresponds to 5 (2*2+1) pixels).

e.g. run : python3 MainAnalysis.py enter.csv results.csv /EmbryoAnalysis /Images /MSC 2


The output 'results.csv' is located in the results/ sub-directory. It contains 7 columns :

1. The embryo name
2. The mean actine level in the all embryo (Actine_all)
3. The mean actine level in cell borders only (Actine_fil)
4. The mean actine level inside cells only (Actine_oufil)
5. The mean bcat level in the all embryo (bcat_all)
6. The mean bcat level in cells borders only (bcat_fil)
7. The mean bcat level inside cells only (bcat_outfil)


------- Step 2. Launch the plotting program

To analyse the different results in the results.csv file, one possibility is to do a boxplot comparing the same signal for different phenotypes. This what does the BoxplotFull.py program in the sub-directory Plot/

It outputs 2 files (one .svg (plot) and one .csv (raw results)) for each of the 6 measures in the results.csv file (e.g. Actine_fil).

In a given plot file, one boxplot is drawn for each phenotype/condition in the enter.csv file. The distributions of the different conditions are compared using a Mann-Whitney two-tailed p-value test.

To launch the program use the following syntax:

python3 Boxplot-Full.py arg1 arg2

arg1 : name of enter.csv file
arg2 : name of results.csv file

e.g. run : python3 Boxplot-Full.py enter.csv results.csv
